# Broad domains of histone marks in the highly compact *Paramecium* macronuclear genome

**DOI:** 10.1101/2021.08.05.454756

**Authors:** Franziska Drews, Abdulrahman Salhab, Sivarajan Karunanithi, Miriam Cheaib, Martin Jung, Marcel H. Schulz, Martin Simon

## Abstract

The unicellular ciliate *Paramecium* contains a large vegetative macronucleus with several unusual characteristics including an extremely high coding density and high polyploidy. As macronculear chromatin is devoid of heterochromatin our study characterizes the functional epigenomic organisation necessary for gene regulation and proper PolII activity. Histone marks (H3K4me3, H3K9ac, H3K27me3) revealed no narrow peaks but broad domains along gene bodies, whereas intergenic regions were devoid of nucleosomes. Our data implicates H3K4me3 levels inside ORFs to be the main factor to associate with gene expression and H3K27me3 appears to occur as a bistable domain with H3K4me3 in plastic genes. Surprisingly, silent and lowly expressed genes show low nucleosome occupancy suggesting that gene inactivation does not involve increased nucleosome occupancy and chromatin condensation. Due to a high occupancy of Pol II along highly expressed ORFs, transcriptional elongation appears to be quite different to other species. This is supported by missing heptameric repeats in the C-terminal domain of Pol II and a divergent elongation system. Our data implies that unoccupied DNA is the default state, whereas gene activation requires nucleosome recruitment together with broad domains of H3K4me3. This could represent a buffer for paused Pol II along ORFs in absence of elongation factors of higher eukaryotes.

## 1 Introduction

The degree of epigenetic differentiation and the organization of eukaryotic genomes is usually adapted to the complexity of an organism: chromatin serves as an additional layer of information, either for manifestation of gene expression patterns, for the cyclic condensation of chromosomes or microtubule assisted separation of DNA in mitotic divisions. Chromatin further influences the proper processing of functional mRNAs as histone modifications influence Pol II dynamics and its interaction with RNA modifying components, such as the capping enzyme or the spliceosome.

*Paramecium tetraurelia* is a unicellular organism belonging to the SAR clade (including stramenophiles, alveolata and rhizaria), which is as distant to plants, fungi, and animals. *Paramecium* is a ciliate, a phylum of alveolatae and shows an unusual nuclear feature: although unicellular, these cells already differentiate between germline and soma by presence of germline micronuclei (Mic) and somatic macronuclei (Mac). Both differ in structural and functional aspects. Micronuclei are small (1-2*µ*m) and transcriptionally inactive during vegetative growth, because the large (approx. 30*µ*m) Mac transcribes all necessary genes to allow for cell proliferation [9]. During sexual reproduction, haploid meiotic nuclei are reciprocally exchanged and fuse to a zygote nucleus: this creates new Mics and Mac while the new developing Mac (anlagen) already transcribes some genes involved in development [23, 52].

The genomic structures between Mic and Mac are quite different. Mics contain thousands of short transposon remnants (IES, internal eliminated sequences), which become deleted by a germline specific RNAi mechanism during macronuclear development [1]. The Mac differs from the Mics by the absence of IESs and transposons [26]. In addition, Mac chromosomes are tiny in size usually below 1Mb, because Mic chromosomes are fragmented into many (∼ 300) different Mac chromosomes. These are amplified then to ∼800 copies each, resulting in a massive polyploidy. Strikingly, the separation of that many DNA molecules (approx. 300 Mac chromosomes x 800n) cannot be handled by classical mitosis. As a result, the Mac divides amitotically: replicated DNA becomes distributed to daughter nuclei without chromosome condensation and without a typical mitotic spindle. The latter would be useless as the absence of centromeres [38] and consequently kinetochors would not allow for attachment of microtubules. However, the amitotic division of Macs in modern ciliates can be seen as a novel feature as e.g. the Karyorelictaea are not able to amitotically divide their Macs; instead they re-generate a Mac each vegetative cell division, meaning every cell cycle [15].

In 2006, the macronuclear genome project revealed two highly unexpected findings: first, an exceptionally high number of genes (∼ 40,000), most of them resulting from three successive whole genome duplications. Second, an exceptionally high coding density of 78%. The latter is due to tiny introns, predominantly of 25bp length, and small intergenic regions (352 bp on average) [5].

Chromatin during amitotical M-phase remains uncondensed suggesting that the Mac does not harbor the full genetic requirements to create highly condensed chromatin. In addition, interphase chromatin was reported to show several unusual features when compared to other species based on chromatin spread preparations. For instance, the finding of several unusual filament types and the appearance of a low level of polyteny between individual transcription nodes [53]. Classical heterochromatin is believed to be absent from the Mac, although a deeper biochemical insight in the Mac chromatin organization is still missing. The same holds true for the presence of classical repressive histone marks in the vegetative Mac, raising the question on how gene repression is regulated. Another epigenetic mark, 5-methylcytosine is known to be involved in negative regulation of gene expression in many eukaryotes. However, 5-methylcytosine is reportedly absent in Mac DNA [57].

Hence, the contribution of dynamic Mac chromatin modifications to the regulation of gene expression remains poorly understood in ciliates. We know from other organisms that chromatin marks have functions in RNA processing and active elongation of transcription. Current studies of mammalian chromatin report functions for well positioned nucleosomes in context of Pol II phosphorylation and interaction with RNA modifying enzymes. This raises the question on how such a regulation is realized in ciliates, specifically in *Paramecium*.

+1 nucleosome positioning, for instance, was indicated to correlate with Pol II pausing and increased recruitment of NELF (negative elongation factor) [33]. Whereas initiation of transcription is accompanied by phosphorylation of serin5, P-TEFb was shown to mediate the conversion of the Pol II complex from its initiation to the processive elongation form, which includes phosphorylation of serin2 [11, 20]. Promoter proximal pausing is known to be controlled by the negative regulators NELF and DSIF, while the C-terminal domain (CTD) of Pol II interacts with the capping components for 5′-capping of the nascent mRNA. Similarly, polyadenylation and splicing are controlled by both, the CTD of Pol II and correctly positioned nucleosomes [10]. Especially for the latter aspect, alternative splicing has been implicated to be regulated by alternative CTD phosphorylation regulated by the SWI/SNF chromatin remodeling complex [7]. Although we do not know much about these mechanisms in ciliates, we suspect them to differ to the above described CTD regulation and interaction with additional components in metazoans. This suspicion arises from the missing Pol II heptameric repeats in *Paramecium*, which likely affect also the interacting complexes due to a co-evolutionary effect. The Mediator complex of *Tetrahymena* for instance, significantly differs to other species [65]. As a consequence, we currently do not understand the role of the ciliate epigenome architecture in relation to Pol II activity in terms of initiation, elongation, pausing and interaction with complexes.

## 2 Materials and Methods

### 2.1 Cell culture and RNA isolation

*Paramecium tetraurelia* cells (strain 51) of serotype A were cultured as described before using *Klebsiella planticola* for regular food in WGP (wheat grass powder) [56]. All cultures for this study were grown at 31°C. To ensure the vegetative state of the Mac, cells were stained with DAPI.

### 2.2 Genomic annotations

The genomic features shown in Figure 2B are captured from the annotations of the respective organisms namely from *Paramecium*DB (strain 51, version 2), *Tetrahymena* Genome Database (version 2014) [59], PomBase (version 2020) [17], and from the ensemble database for *Drosophila melanogaster* (release 98), and *Homo sapiens* (release 100) [64].

**Figure 1.**
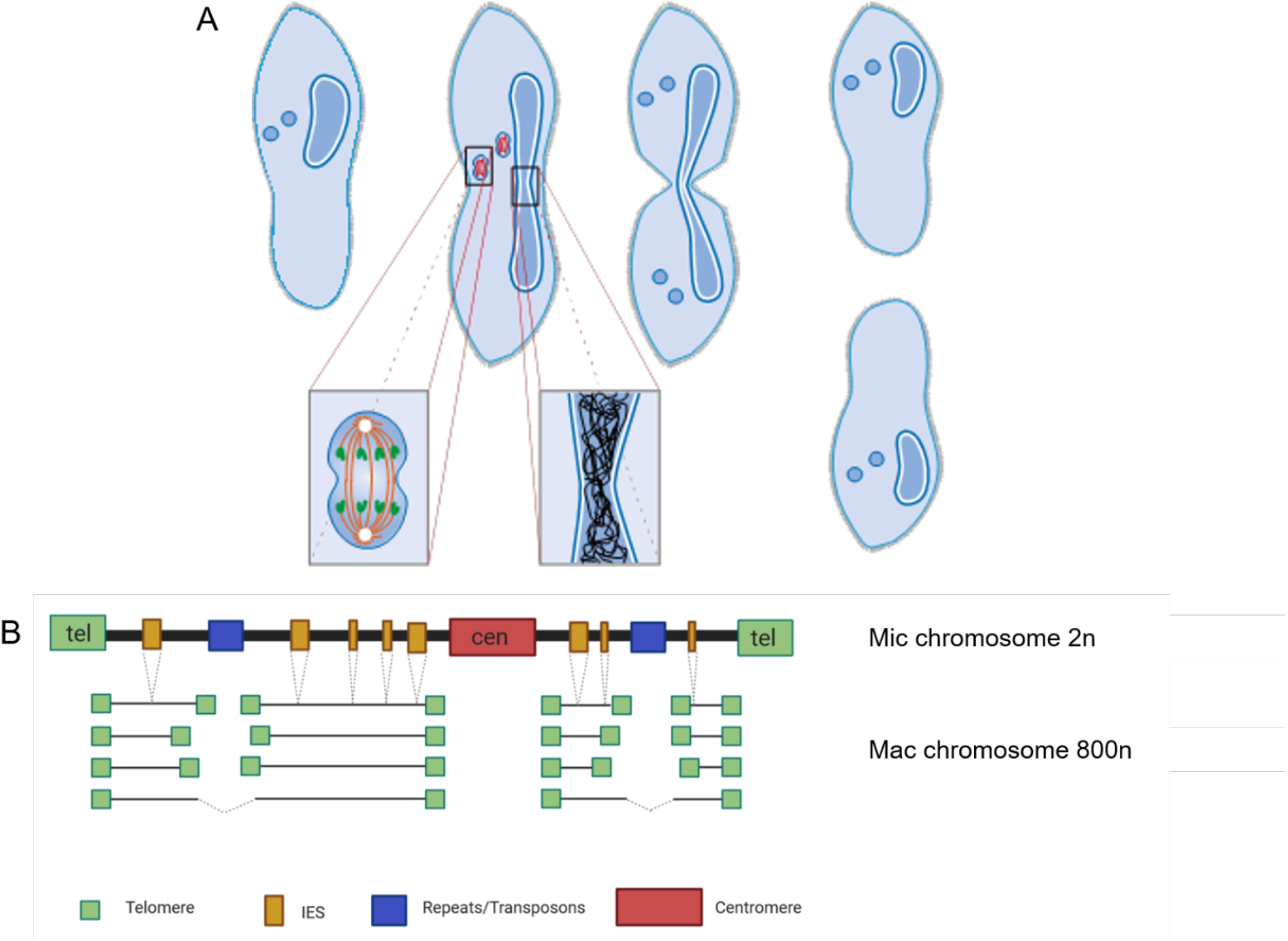
*Paramecium* vegetative cell divisions and chromosomal structure of Mic and Mac. A, *Paramecium tetraurelia* showing two generative Mics and one vegetative Mac. Cell division involves mitotic separation of condensed Mic chromosomes and amitotic separation of uncondensed Mac chromosomes. While Mics and Mac divide the nuclear envelope remains at both nuclei. (Figure courtesy of Jens Boenigk and Martin Simon) B, Chromosomes of the diploid Mic are large and contain centromeres and telomeres similar to canonical eukaryotic chromosomes. In addition, they consist of ∼60.000 IES elements (internal eliminated sequences) and repeats (transposons, minisatellites). During macronuclear development after sexual reproduction (not shown here), telomeres, centromeres, repeats and IES become eliminated by different mechanisms. While IES are precisely excised, elimination of repeats and presumably centromeres occurs imprecise resulting in fragmentation into heterogenous macronuclear chromosomes (with rare fusion of fragments). All macronuclear fragments show *de novo* telomere addition and amplification to 800n. (Created with BioRender.com)

**Figure 2.**
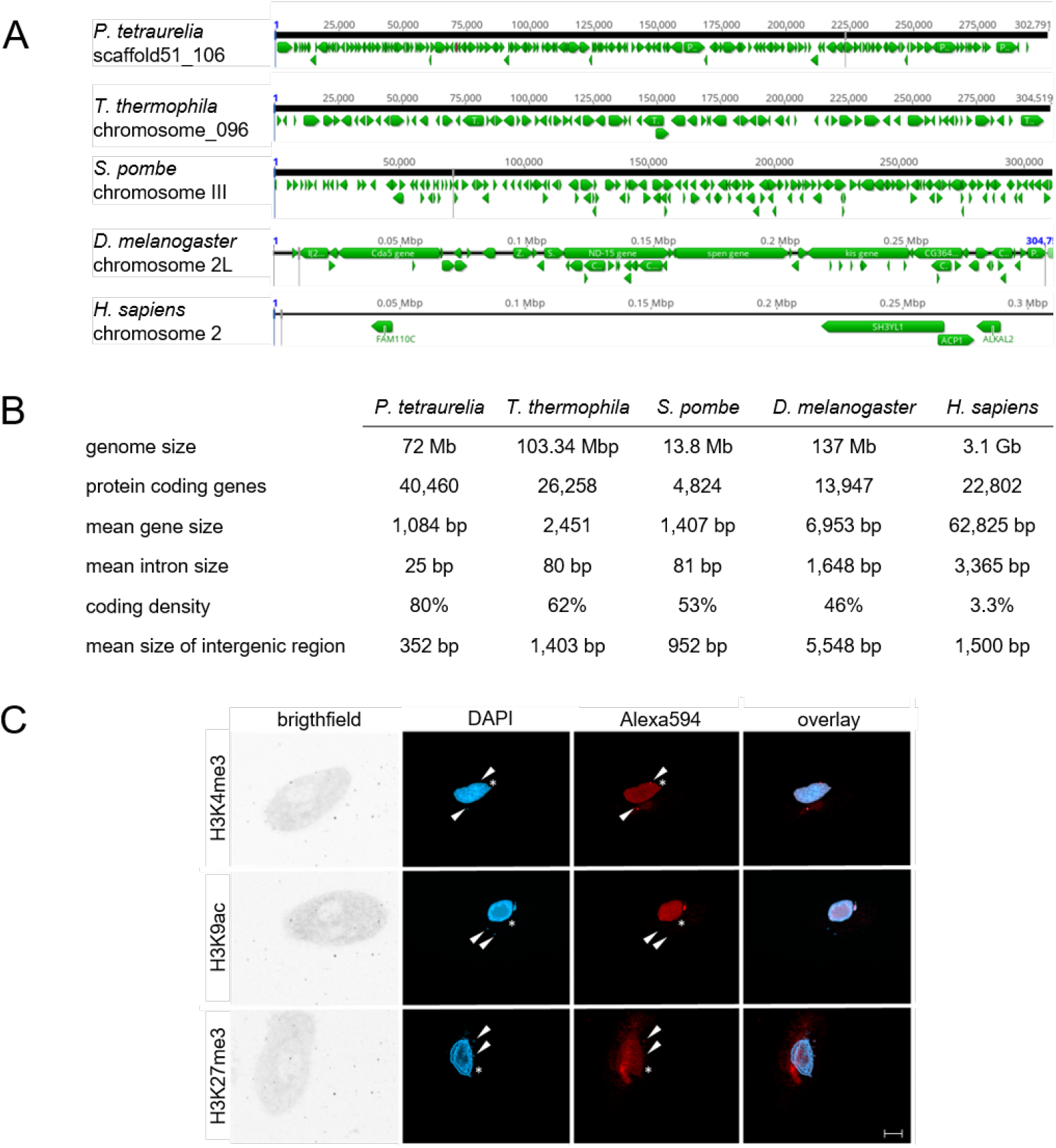
Features of the *Paramecium* genome in comparison to other organisms. A, Comparisons of distribution of genes (green arrows) along the chromosomes of selected organisms to highlight the variation in coding density (*Paramecium tetraurelia, Tetrahymena thermophila, Schizosaccharomyces pombe, Drosophila menlanogaster, Homo sapiens*). A window of 300 kb is shown for each chromosome in a genome browser. B, Summary of genomic features of the same organisms named in A. See material and Methods for details on collected data. C) Detection of histone modifications in vegetative *Paramecium* nuclei by immunofluorescence staining. DNA in the nuclei is stained with DAPI (blue) while antibodies directed against the three indicated modifications (H3K4me3, H3K9ac, H3K27me3) were labeled with a secondary Alexa594 conjugated antibody (red). Arrowheads point at micronuclei, asterisks indicate position of the macronucleus. Other panels show brightfield and overlay of signals. Representative overlays of Z-stacks of magnified views are shown. Scale bar 10 *µ*m.

### 2.3 Antibodies

Polyclonal, ChIP-seq grade antibodies directed against histone modifications were purchased from Diagenode: H3K9ac # C15410004, H3K27me3 # C15410195, H3K4me3 #C15410003. For antibody against *P. tetraurelia* RBP1, the peptide SPHYTSHTNSPSPSYRSS-C was used for rabbit immunisation. Purification and testing of specificity by Western blots and immunostaining was carried out as described recently [18]. Since there are some amino acid differences in the N-terminal tail of the *Paramecium* H3P1 to *Human* H3 (Suppl. Fig.1A), the peptide PtH3K27me3 TKAARK(me3)TAPAVG was synthesized and binding affinity of the purchased H3K27me3 antibody to the PtH3k27me3 peptide was verified by dotblots and competition assays. Peptide competition assays (Suppl.Fig.1B) were performed by blocking 2 *µ*g of each antibody with a 10 fold excess of its corresponding peptide over night at 4 °C with agitation. 1 pmol to 100 pmol of each peptide were blotted on a nitrocellulose membrane and decorated with blocked and unblocked antibodies.

### 2.4 Fixation of cells

Isolation of intact macronuclei from fixed cells was carried out using an adapted NEXSON protocol [4]). 2-3 million cells were washed twice in Volvic® and starved for 20 min at 31 °C. After harvesting (2500 rpm, 2 min), the cell pellet without remaining media was resuspended in 2 ml fixative solution (20 mM Tris-HCL pH 8, 0.5 mM EGTA, 1 mM EDTA, 10 mM NaCl, 1 % methanol-free formaldehyde). After incubation (15 min, room temperature), the reaction was quenched by adding glycine to a final concentration of 125 mM. Cells were centrifuged (3300 g, 3 min, 4 °C) and the supernatant was discarded. The pellet was washed once in ice cold PBS buffer and once in PBS buffer supplemented with cOmplete Protease Inhibitor Cocktail, EDTA-free (PIC, Roche, #11873580001). Cell suspension was split in half, centrifuged (3300 g, 5 min, 4 °C) and cell pellets were flash frozen in liquid nitrogen.

### 2.5 MNase-seq

One aliquot of cell pellet was thawed on ice, re-suspended in 2 ml Farnham lab buffer (5 mM PIPES pH 8, 85 mM KCl, 0.5 % NP-40) and evenly split into pre-cooled 1.5 ml Bioruptor tubes (Diagenode). After sonication (15 sec on/ 30 sec off, 5 cycles, 4 °C) using Bioruptor 300 (Diagenode) 5 *µ*l were stained with DAPI to verify isolation of intact Macs. Cell suspension was centrifuged twice (3000 g, 5 min, 4 °C) with washing of the pellet in Farnham lab buffer in between. The following isolation of DNA covered by mononucleosomes was isolated as described in [63]. One aliquot of isolated nuceli was resuspended in 1x MNase buffer (50 mM Tris-HCL pH 8.0, 5 mM CaCl_2_) and split into portions of 20.000 nuclei per reaction. After centrifugation (3000 g, 5 min, 4 °C) nuceli pellets were re-supended in 500 *µ*l MNase reaction buffer (50 mM Tris-HCl pH 8.0, 5 mM CaCl_2_, 10 mM *β*-Mercaptoethanol, 1% NP-40, 500 ng BSA). To each reaction, 10 or 128 units of MNase (NEB, # M0247S) was added and after incubation (10 min, 37 ° C, 450 rpm), the reaction was stopped (10 mM EGTA, 1 mM EDTA, 5 min, 450 rpm). DNA corresponding to the size of mononucleosomes (100-200 bp) was re-isolated from a 3% agarose gel using MinElute Gel Extraction Kit (Qiagen, # 28604). As input, nuclei were treated with Proteinase K, extracted as described and treated with 0.1 U or 1.5 U MNase (5 min, 28 °C) and extracted again. DNA was load onto a 3% agarose gel and mononucleosomal fractions (100-200 bp) were re-isolated. DNA library preparation was performed using NEBNext® Ultra™ DNA Library Prep Kit for Illumina® (NEB, E7370) with 10 ng input, 11 PCR cycles and KAPA Taq HotStart DNA polymerase (Kapa Biosystems, # KK1512). MNase-seq read count correlation of four independent replicates, each, used for subsequent analyses can be found in Suppl. Fig.2 as well as a comparison of nucleosome occupancy resulting from 10U (light) and 128U (heavy) digestions.

### 2.6 Chromatin immunoprecipitation (ChIP-seq)

Nuclei pellets originating from the same fixed cells as used for MNase treatment were re-suspended in shearing buffer (10 mM Tris-HCl pH 8, 0.1% SDS, 1 mM EDTA) and transferred in fresh, pre-cooled Bioruptor tubes. For shearing of chromatin, suspension was sonicated (30 sec on/ 30 sec off, 5 cycles, 4 °C). After centrifugation (16000 g, 10 min, 4 °C) the supernatant containing the chromatin was aliquoted in 100 *µ*l portions and stored at −80 °C. To control shearing efficiency, 50 *µ*l of each chromatin aliquot were de-crosslinked using Proteinase K (20 mg/ml), followed by phenol/chloroform/isoamylalcohol extraction, which was repeated after RNase A (10 mg/ml) digestion. DNA was precipitated and concentration was measured using NanoDrop. Aliquots of 2 *µ*g were run on a 1.5% agarose gel. 8 *µ*g of adequately sheared chromatin was subjected to immunoprecipitation using iDeal ChIP-seq kit for Histones (Diagenode, C01010050) with 2 *µ*g of antibodies against histone modifications or 10 *µ*g of custom RPB1 antibody. Input was generated by putting 1 *µ*l of chromatin aside without mixing to antibodies. After overnight IP and elution from the magnetic beads, precipitated chromatin was de-crosslinked, RNase A treated and extracted as described above. DNA library preparation was performed using NEBNext® Ultra™ DNA Library Prep Kit for Illumina® (NEB, #E7370) with 10 ng input, 11 PCR cycles and KAPA Taq HotStart DNA polymerase (Kapa Biosystems, # KK1512).

ChIP-seq read count correlation of four independent replicates of H3K4me3, H3K27me3, H3K9ac IP each, used for subsequent analyses can be found in Suppl. Fig.3.

### 2.7 Sequencing and pre-processing

Prepared libraries were quantified using the dsDNA HS assay for Invitrogen Qubit 2.0 Fluorometer (ThermoFisher) and size distribution was measured with the Bioanalyzer High Sensitivity DNA Kit (Agilent). DNA libraries resulting from MNase digestion and ChIP were sequenced on an Illumina HiSeq2500 in high output run mode. All histone ChIP-seq reads were first trimmed for adapter sequence and low quality tails (*Q <* 20) with Trim Galore (v.0.4.2) [35, 42]. We utilized deeptools2 [49] to investigate the quality of replicates (*multiBamSummary, plotFingerprint* and *plotCorrelation* tools) with subsequent down sampling of some histone ChIP replicates, which had rather high coverage (see Suppl. Table 1; sheet sequencing depth). All raw read data of this study has been deposited at ENA, accession no. PRJEB46233.

### 2.8 Alignments

All MNase, Pol II and histone ChIP-seq reads were aligned to the macronuclear reference genome *P. tetraurelia* (strain 51, version 2) [2] after quality control. Alignments were performed using the local mode of bowtie2 [36] software with default parameters except the seed alignment mismatch parameter which was set to 1 (-N 1). We used these alignments for the subsequent steps described in sections 2.10 and 2.12. For the steps described in 2.11, we used histone ChIP-seq alignments performed using the default parameters of the GEM mapper [41] and then duplicated reads were annotated with Picard tools (v1.115) (http://broadinstitute.github.io/picard). Amino acid sequences from RPB1 subunits were aligned by ClustalW and visualized in Geneious Prime 2020.2.2 and BioEdit [27] (Accession Nos.: H.s. P24928; S.p. NM001021568; S.c. YDL140C; T.t. 00538940; P.t. PTET.51.1.P1370127).

### 2.9 Expression and intron data

We utilized the mRNA expression data of strain 51 wildtype serotype A from our previous work [12] which can be accessed at European Nucleotide Archive (ENA) with the accession PRJEB9464. We quantified the expression using Salmon (v0.8.2) [47] default parameters for all replicates, and utilized the mean of replicates in all downstream analyses. We used the transcript annotation from the MAC genome of *P. tetraurelia* (version 2; strain 51 [3]). For creating intron profiles, we created a 20 bp window centred on the first and last intron base of the 5’-exon-intron junction and the 3’-intron-exon junction. We plotted the nucleosome profile for 1500 bp around this window with the centre of x-axis representing the junctions (see Figure 4C).

**Figure 3.**
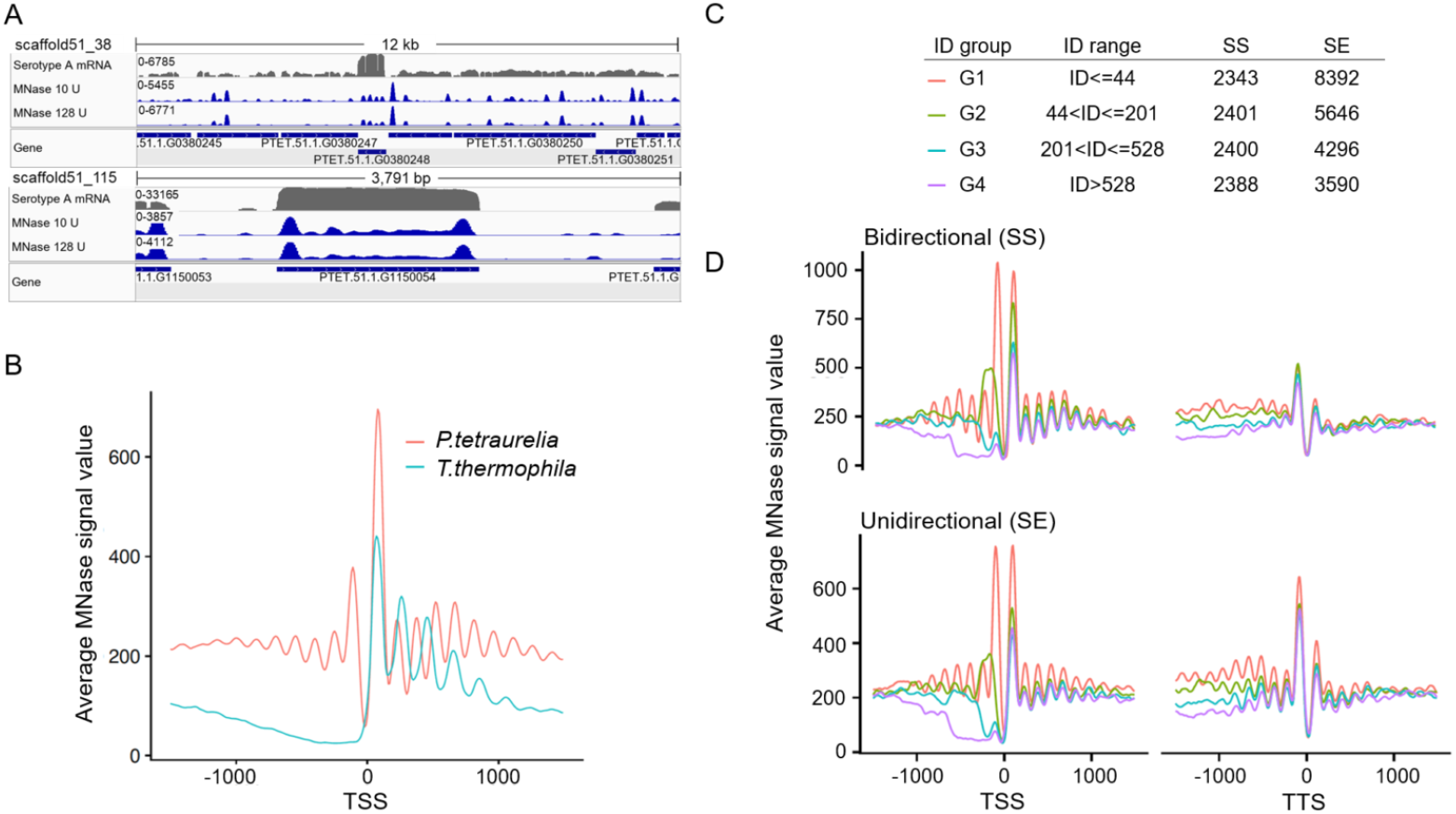
MNase - seq results reveal well positioned +1 nucleosomes. A, Exemplary view of nucleosome distribution along the Mac scaffolds of *Paramecium*. Top panel shows the peak distribution in a 12 kb window while the lower panel shows the magnified view on one gene. For both panels, the top row shows the coverage track from polyA mRNA - seq followed by the tracks for nucleosome occupancy obtained by light (10U) and heavy (128U) MNase digestion of *Paramecium* nuclei. Coverage tracks were visualized using IGV browser [50]. B, Profile plot for nucleosome distribution in relative distance to the transcription start site (TSS) for all analyzed Paramecium genes. Signal for 1000 bp up- and downstream of the TSS are shown. For comparison, MNase - seq data from *T*.*thermophila* was plotted in the same manner. C, Dissection of neighbouring *Paramecium* genes based on their configuration and intergenic distance (ID). Table shows separation of genes by configuration and ID, ranked from short distances (G1) to long distances (G4). The last two columns indicate numbers of genes in each configuration and ID group. D, Nucleosome profiles in a 2KB window centered at the TSS (left) or the transcription termination site (TTS, right) for neighbouring genes in SS and SE configuration are shown. Genes were additionally separated by the length of their intergenic distances colour coding in C.

**Figure 4.**
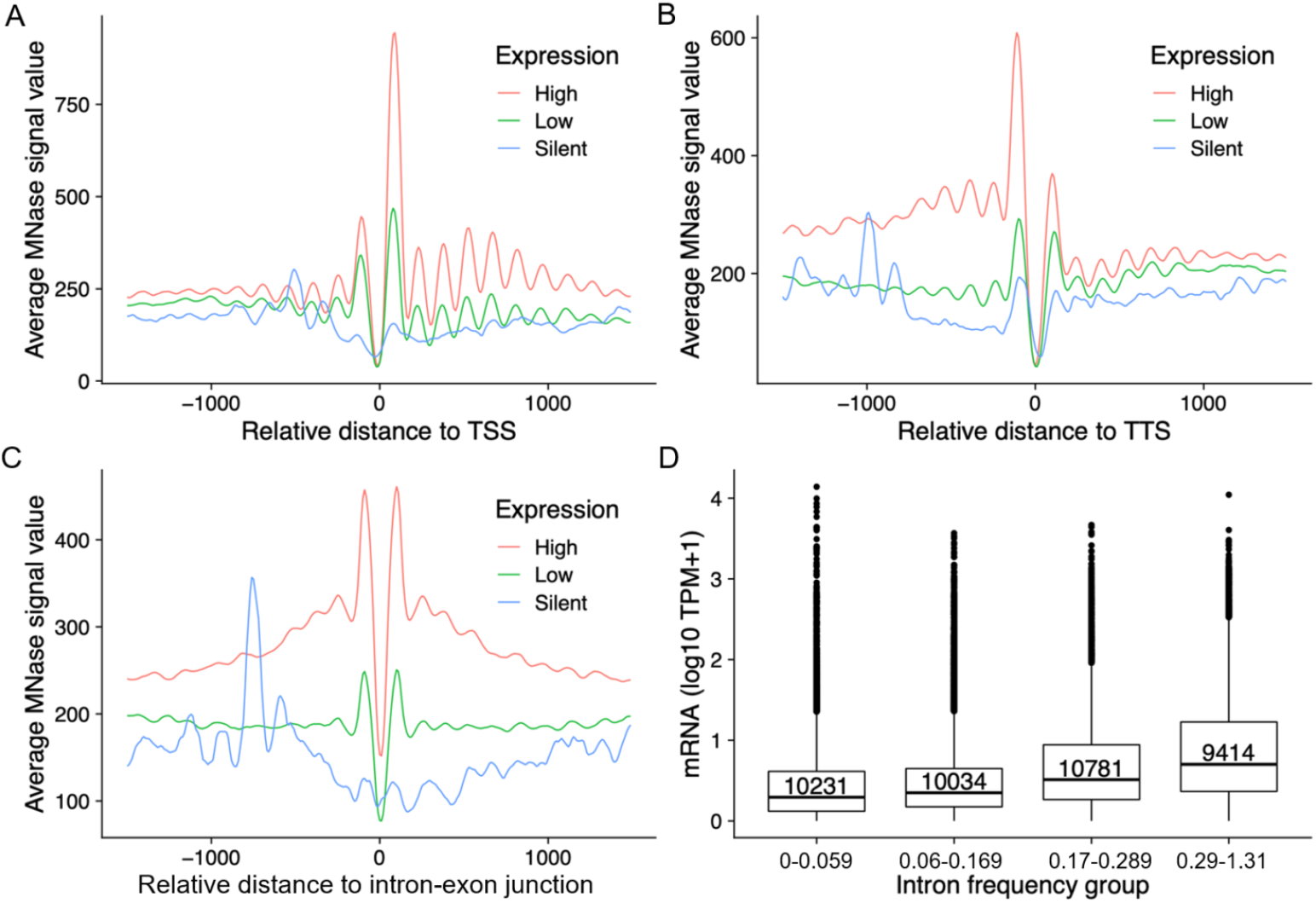
Positioning of nucleosomes in correlation to gene expression. The nucleosome profiles in relation to their distance (x-axis) to TSS (A), TTS (B), and intron-exon junction (C) is shown for gene categories based on their expression levels. D) Box plots showing the mRNA expression (y-axis; log10 TPM+1) of genes with different intron frequency groups (number of introns per 100 bp; x-axis). A Kruskal-Wallis test showed that the expression distribution between all pairs of intron frequency groups is significantly different (P < 2.2e-16).

### 2.10 Peak calling

We used the DANPOS2 [14] software to perform position or peak calling. We used the *dpos* functionality to call the positions of MNase and Pol II peaks and the *dpeak* functionality for histone ChIP peak calling. Default parameters were used for all functionalities of DANPOS2. Further, we made use of the *profile* functionality of DANPOS2 to visualise how a chromatin feature is distributed in a genomic annotation of interest (See Fig.3).

### 2.11 Segmentation analysis of chromatin marks

We employed ChromHMM [21] to perform genome-wide segmentation using the histone marks (H3K27me3, H3K4me3, H3K9ac), and MNase data. First, the genome was binarized into 200bp bins based on a Poisson background model using the *BinarizeBam* function. Second, the binarized data was used to learn a chromatin state model with 5 states using the *LearnModel* function. The states were then annotated to different genomic annotations. We used the *plotProfile* and *plotHeatmap* functionality of deeptools2 to create scaled enrichment plots of different chromatin features. In this context, scaling refers to shrinking or stretching a genomic locus to a fixed length set by user. In the plots we have often scaled the loci in between transcription start site (TSS) and transcription termination site (TTS) to 1500 bp unless mentioned otherwise in the figures.

### 2.12 Comparative Pol II analysis and pausing index

We used the data sets mentioned in Supplementary Table 1 for the comparative Pol II analysis of different organisms shown in Figure 6. We calculated the pausing index, after applying a threshold on the number of reads in the TSS Region of genes (see 2.12), depending on the distribution of read counts of individual data sets. The thresholds are mentioned in Figure 6C. The mRNA quantification was done using default parameters of Salmon with transcripts obtained from the respective genomic annotations mentioned above. The mean of replicates were used in all cases. We defined a region starting at 30 bp upstream of TSS till 300 bp downstream of TSS as *TSS region*, and a region starting at 300 bp downstream of the TSS until the TTS as *gene body*. The pausing index is calculated as a ratio of reads (in TPM) in the TSS region compared to reads in the gene body. Genes with a pausing index greater than 1.5 were considered as paused.

**Figure 5.**
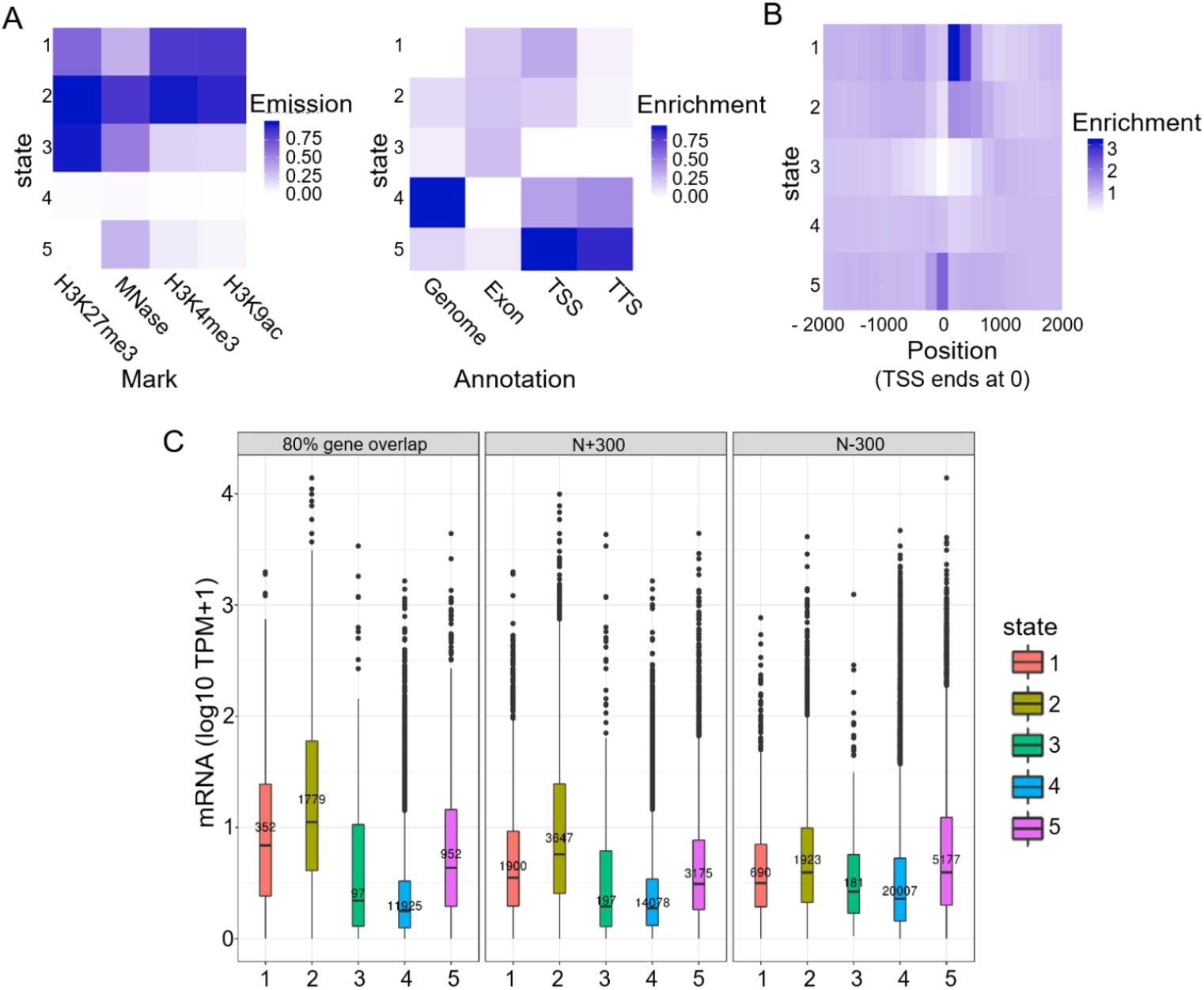
Segmentation analysis using ChromHMM. A, The chromatin state assignments are shown as a heatmap of emission parameters from a 5-state ChromHMM model (left). Each row corresponds to a ChromHMM state, and each column represents a different epigenetic mark. The darker the color of an epigenetic mark for a state the higher the probability of observing that epigenetic mark in that state. Heatmap showing the overlap fold enrichment of each ChromHMM state (row) in different genomic annotations (columns, right). Enrichment values are obtained from the overlap enrichment functionality of ChromHMM with a column-specific color scale. B, The fold enrichment of each state in 200 bp bins within a 2 kb window around the transcription start site (TSS) is shown. Enrichment values are obtained from the neighborhood enrichment functionality of ChromHMM with a uniform color scale. C, Box plots showing the mRNA expression (y-axis; log10 TPM+1) of genes whose loci overlap at least by 80% with a respective state (left). Additionally, genes were separated by their assigned state at the first 300 bp of the gene body (N+300) and 300 bp upstream of the TSS (N-300) and mRNA expression values of these genes is plotted (middle, right).

**Figure 6.**
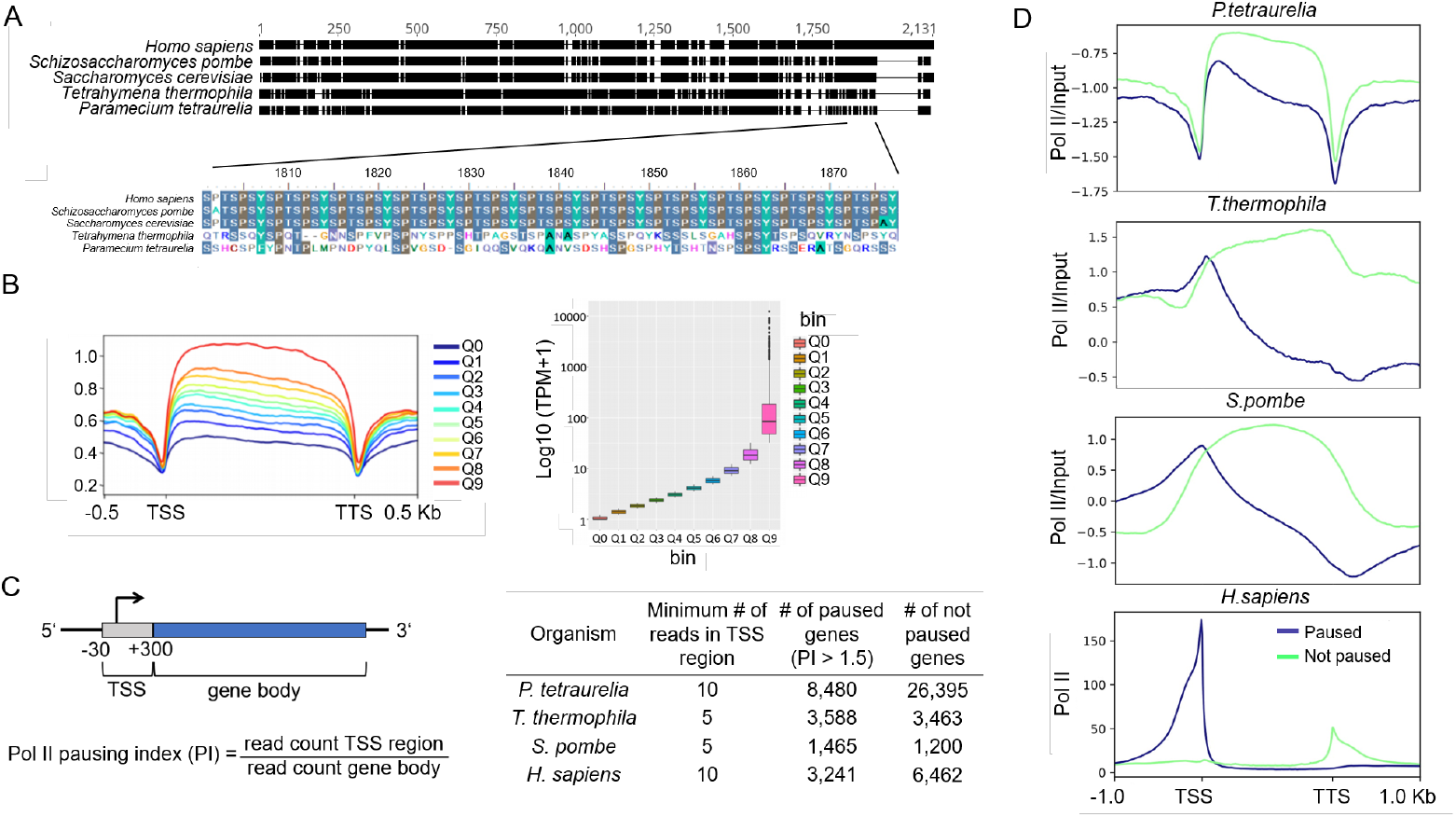
Analysis of RNA polymerase II pausing. A, Multiple sequence alignment of the RNA polymerase II enzyme’s RPB1 subunit in different organisms is shown. The C-terminal end of RPB1 is zoomed in to show the difference in conserved regions of some ciliates to other organisms. B, (right) Box plots of gene expression (y-axis; log10 TPM) split in 10 quantiles is shown; higher quantiles means higher expression. (left) Pol II enrichment (y-axis) profiles of genes in respective quantiles are shown. Distance shown on the x-axis is scaled, i.e. all genes (TSS-TTS) are either stretched or shrunken to a length of 500 bp. A 500 bp window up- and downstream of the gene loci is included. Enrichment profiles were plotted using deeptools2. C, A graphical representation of the regions included in polymerase pausing index (PI) calculation is shown. We categorized a gene as paused if the PI ≥ 1.5. The table summarizes numbers of paused/not paused genes for selected organisms (Suppl. Tab. 1 contains details on Pol II datasets). D, Same as the Pol II enrichment profiles in B but genes are split based on the status of Pol II pausing.

### 2.13 Classification of gene expression using random forests

After removing 1369 silent genes whose mean expression is zero (*TPM* = 0), we split the remaining genes into 19,090 high (*TPM* > 2) and 20,001 low expressed genes (*TPM <* 2). Cut-offs were determined using the first quartile of the distribution of wildtype 51A sertoypes mRNA expression. For these gene sets, gene body normalized read counts were calculated of H3K27me3, H3K4me3, H3K9ac, Pol II, and MNase, called epigenetic features and in addition, the ratio of H3K4me3 and H3K27me3. We also obtained three genetic features: gene length, intron frequency, and intergenic length. Using these features and the labels (high/low expressed), we built a random forests classifier in python (version 3) using the default parameters available with the scikit-learn package [48]. As our intention is to understand the relation between expression and these different features, we used all available data to train the model using a 40-fold cross validation (CV) method. We use the CV based area under the precision-recall curve (PR-AUC) to evaluate the performance of different models. A PR-AUC of 1 would represent a perfect model, which 100% of the times would correctly predict whether a gene is highly or lowly expressed. Further, we used the shap package [39] to calculate the global and local feature importance.

### 2.14 Partial correlation networks

We investigated the partial correlation of any two epigenetic marks of interest, after removing the effects of other measured epigenetic marks by using the sparse partial correlation networks method [37]. We used the gene body normalized signals of all the epigenetic marks in this study, and the mRNA expression for this analysis.

### 2.15 Analyses of gene expression plasticity

Using the available transcriptome data [12], plasticity of genes was calculated step wise: First, the mean TPM for each gene over different conditions (expression data from Serotype A, B, D, H as well as heat shock conditions) was calculated. To see if the gene expression is fluctuating or stable around the mean value, the absolute deviation from the mean for each gene was calculated. The higher this value, ranging from 0.07 to 1.79, the higher is the fluctuation in gene expression. We refer to genes with a large fluctuation as *plastic genes*. For the random forests analysis of plastic genes, we grouped all genes in four groups of roughly similar gene numbers. Then we performed random down-sampling of highly or lowly expressed genes such that there is an equal number of genes in both groups for classification (sub-samplng done five times).

## 3 RESULTS

### 3.1 Unusual properties of the macronuclear genome

In this work, we aim to understand the epigenomic organisation of the polyploid vegetative Mac of *Paramecium tetraurelia*. These cells contain two diploid and transcriptionally silent micronuclei, which divide by classical mitosis during cellular fission, while the mac divides amitotically: stretching and outlining results in uncontrolled separation of uncondensed chromosomes (Fig.1A). Interpretation of any Mac epigenome data requires a look for the genomic structure of the chromosomes. During their processing from Mic chromosomes after sexual recombination, heterochromatic regions such as telomeres, centromeres, satellites and transposons become eliminated in addition to ∼60.000 transposon remnants called internal eliminated sequence (IES) elements (Fig.1B). Fragments undergo *de novo* telomere addition resulting in small acentromeric chromosomes with a size below 1MB. These chromosomes exist at varying lengths due to imprecise eliminations of repeated sequences [19]. Compared to other species, even the related ciliate *Tetrahymena*, the *Paramecium* Mac genome shows an extremely high coding density of about 80% with small intergenic regions and tiny introns of 25nt [5]. These features become even more striking in comparison to *S*.*pombe* and individual metazoens (Fig.2A/B).

In order to quantify global epigenome organisation in *Paramecium*, we first investigated the distribution of histone H3 modifications in the vegetative Mac, since histone modifications are major contributors to chromatin architecture. Immune fluorescence analysis with histone H3 specific Abs show H3K4me3 and H3K27me3 occurring in both, Mics and Mac, while H3K9ac is present in the Mac, only (Fig.2C).

### 3.2 Well positioned nucleosomes locate at TSS

To characterize nucleosome positioning, mono-nucleosomal DNA was isolated after digestion of macronuclear chromatin with micrococcal nuclease (MNase). Reads were mapped to the genome assembly resulting in discrete peaks for both setups using 10 or 128U MNase (Fig.3A), corresponding to light and heavy digestion. Genomic analysis of MNase data revealed well positioned +1 and −1 nucleosomes at the transcription start site (TSS) (Fig.3B). Especially the presence of −1 nucleosomes differs to analog analyses of MNase data from *Tetrahymena, S*.*pombe, D*.*melanogaster*, but they are apparent in humans (Suppl. Fig.4). As such their presence in *Paramecium* is surprising and requires additional analysis. In addition the comparison to other species shows that downstream nucleosomes (downstream of +1) in *Paramecium* are apparently much less pronounced, already the +2 nucleosome signal is roughly background, which is in contrast to *Tetrahymena, S*.*pombe* and *Drosophila* showing slightly decreasing peak values inside the gene bodies (Suppl. Fig. 4).

In the following, we aimed to see whether the positioning of −1 nucleosomes could be due to short intergenic regions. We therefore dissected the *Paramecium* genes due to two parameters: intergenic distance and orientation of genes. We considered bidirectional promoter genes, where the two start sites of both genes are adjacent (Start-Start,SS), or unidirectional genes where one start site is paired with the end of the other gene (Start-End, SE, Suppl. Fig.5A). These two categories were additionally classified into four groups based on their intergenic distance. The number of genes in each category is given in Fig.3C. Fig.3D shows nucleosome positioning of these categories at the transcription start site (TSS) and the transcription termination site (TTS). Most apparent, putative −1 nucleosomes are much more pronounced in genes with short 5′-intergenic regions below 142bp and this is true for the SE and the SS configuration. In addition, the TTS also shows well positioned nucleosomes at the ultimate 3′-end of ORFs, and these are more pronounced in the SE configuration.

Absence of −1 nucleosomes in genes with longer intergenic region let us conclude that these are either +1 or TTS nucleosomes of upstream genes, but no true −1 nucleosomes. Interestingly, they are still in perfect phasing with the gene of interest. One may hypothesize that *cis* factors regulating nucleosome positioning also control gene distance to synchronize phasing of close genes.

We consequently asked for a potential co-regulation of genes at bidirectional promoters. Correlation analysis of neighboring genes suggest a high degree of co-regulation of all neighbor genes regardless of the configuration (Suppl. Fig.5A/B). However, Suppl. Fig.5C shows that we cannot identify a higher degree of co-regulation in genes under the same bidirectional promoter suggesting that even short intergenic distances are sufficient to control regulation of gene expression independent of the neighbor gene. However, our data indicates that genes with bidirectional promoters tend to a longer intergenic distance (Suppl. Fig.5D) suggesting that selection pressure acts on these regions to separate bidirectional genes from each other. Gene length itself seems not to have a strong effect on TSS and TTS nucleosome positioning (Suppl. Fig.6).

### 3.3 Low nucleosome occupancy at silent genes

We sought to investigate whether nucleosome positioning is changed with differences in gene expression levels (Fig.4A and B). At both ends of a gene, TSS and TTS, well positioned nucleosomes can be found in highly expressed genes only. In contrast, these regions and also gene bodies of silent genes appear to be almost devoid of well positioned nucleosomes.

We can detect well positioned di-nucleosomes around introns (Fig.4C). As mentioned, the 25nt introns are among the shortest reported in eukaryotes [51]. Intron splicing appears to result from efficient intron definition, rather than exon definition as in multi-cellular species, although only three nucleotides define the 5′- and 3′-splice sites [32]. Our data does not reveal any associations of intron nucleosomes with intron length (Suppl. Fig.7A). As our MNase data suggests a general low occupancy of nucleosomes in gene bodies, intron associated di-nucleosomes could be an exception to this. We correlated the intron frequency (number of introns per 100bp) with gene expression levels (Fig.4D) and found increasing mRNA levels with increasing intron frequency, an effect that is independent of the gene length (Suppl. Fig.7B). Thus, introns in *Paramecium* may be involved in transcriptional regulation by recruitment of nucleosomes to gene bodies.

### 3.4 Broad histone mark domains in gene bodies

In order to extent the chromatin analysis to histone modifications, chromatin immuno-precipitation followed by sequencing (ChIP-seq) was carried out from vegetative cells. We used the NEXSON procedure [4] involving isolation of intact Macs without Mics.

Another advantage of this procedure was that we were able to use the very same Mac preparations for both, MNase- and ChIP-seq. We used antibodies for the activation associated marks H3K9ac and H3K4me3, as well as an antibody for the repressive mark H3K27me3. The observed ChIP-seq signatures of these three marks showed rather broad signals, which were not comparable to sharp peaks of metazoen ChIP-Seq signals. Thus, we refrained from a peak-calling approach and used ChromHMM [21] to segment the entire Mac genome into 200bp bins for *de novo* determination of re-occurring combinatorial and spatial signal patterns. We found that five different stable chromatin states could be observed (trying to increase the number of states resulted in highly similar states and we therefore continued all further analyses with five states). Heatmaps in Fig.5A show the contribution of the individual signals to each chromatin state (CS) and on the right, the quantitative assignment of each chromatin state to different regions of the genome. We abbreviate all five chromatin states as CS1 to CS5.

One major finding of the segmentation is represented in CS4. ChromHMM defines this state as being almost free of any signal, this state is moreover attributed to the highest percentage of the genome (Fig.5A, right). This may support our previous assumption, that a high amount of Mac DNA is free of nucleosomes and therefore also of transcription altering histone marks. In contrast, MNase and histone mark signals can be found in CS1-CS3 and CS5. Their ChromHMM signature shows dynamic combinations between the three investigated histone marks and the occurrence of these states also varies in different genomic areas. Focusing on histone marks around the TSS, CS1 and CS2, both enriched in H3K9ac and H3K4me3, show strong accumulation at the +1 nucleosome (Fig.5B). All other chromatin states show derichment at +1, especially CS3, which suggests that especially H3K27me3 is depleted at these gene loci.

To go deeper into the role of the individual marks and states in association with gene expression, we dissected genes into categories overlapping with a chromatin state (i) for more than 80% of the entire gene body, (ii) with first 300bp of the ORF or (iii) with 300bp of the non-coding upstream region. We consequently correlated this with the gene expression level of these genes (Fig.5C). Genes with high levels of H3K9ac and H3K4me3 (CS1) are highly expressed. Focusing to the role of H3K27me3 its high abundance in CS2, associated genes showing the highest expression level, is an argument against a repressive function of this histone mark. Only few genes (91) can be attributed to CS3, the only state where the H3K27me3 signal dominates over H3K4me3 and H3K9ac; although the genes appear to be quite low expressed the small number of genes does not allow for a conclusion about a possible repressive function of H3K27me3.

Genes associated with CS5 show low levels of H3K4me3 and H3K9ac with absence of H3K27me3 and these genes show an intermediate gene expression level. CS4 shows apparently the lowest gene expression level and, in agreement with the quantitative analysis, the highest number of genes. We conclude that gene silencing in the Mac is associated with genomic loci which consist predominantly of free and accessible DNA. Comparing the 80 % gene overlap category to the upstream and the 5′-coding region, our analysis indicates that the upstream region contributes less to gene regulation. Mainly the 5′-CDS and the ORF appear to be involved in gene regulation, which fits to our conclusions from MNase data. We can therefore conclude that gene transcription is mainly associated with high levels of H3K9ac and H3K4me3 at the +1 nucleosome. We do not see a direct evidence for a repressive function of H3K27me3. These results now raise several questions, especially about the role of the prominent +1 nucleosome in transcriptional activation: could this be a place for RNA Polymerase II pausing in order to regulate gene expression?

### 3.5 Pol II occupancy correlates with gene expression levels

In order to characterize Pol II occupancy and activity, it is important to note that *Paramecium* Pol II diverges from conserved metazoen and most unicellular Pol II. In *Paramecium*, as well in *Tetrahymena*, the consensus serine rich heptad repeats are missing, but the CTD shows overall a high percentage of serines (Fig.6A). As commercial Pol II antibodies target the heptamers in the CTD we had to produce an own antibody against the *P*.*tetraurelia* CTD of RBP1. After affinity purification and specificity checks by IF and Western blots of cellular fractions (Suppl. Fig.8), ChIP was carried out as described. Figure 6B shows high Pol II occupancy genes showing high expression and vice versa. Here, the analysis of all genes of the genome results in a quite equal distribution of Pol II along the ORF.

We consequently asked whether Pol II pausing at the +1 nucleosome can be observed and calculated a pausing index (PI) by dividing the Pol II coverage of the TSS by the coverage of the gene body (Fig.6C). Dissecting paused and non-paused genes by a threshold of PI larger than 1.5, we compared Pol II occupancy of *Paramecium* to other species. Fig.6D shows that *Paramecium* is the only species with convergent occupancy of paused and non paused genes. The overall distribution of *Paramecium* Pol II is highly different to other species. In human, *S*.*pombe* and *Tetrahymena*, non-paused genes show increasing coverage along the ORF (see Suppl. Fig.9A for detailed heatmaps). This is different in *Paramecium*, where non-paused genes show in general higher occupancy and less decrease along the ORF. The pattern of *Paramecium* appears different to other species, suggesting that regulated pausing at the +1 nucleosome occurs only rarely. This is to some extent also true for *Tetrahymena* and yeast with the difference that paused genes here show a clearer peak at the TSS along with a strong decrease along the ORF. Such patterns cannot be identified in *Paramecium. Paramecium* in contrast shows a clear drop in Pol II occupancy before the TSS and at the TTS: this seems in agreement with our hypothesis, regulation of gene expression occurs mainly inside ORFs. We further analyzed whether pausing is associated with reduced full length mRNA production. Suppl. Fig.9B shows that we see a significantly lower expression of paused genes in *Tetrahymena* and *S*.*pombe*; only in humans, paused genes show higher mRNA levels. Thus, PolII pausing may indeed be a mechanism of gene regulation, but used in a totally different manner in lower eukaryotes and mammals. Especially in *Paramecium* the mRNA levels between paused and non-paused genes show the smallest differences, although significant: suggesting that pausing is more involved in fine tuning transcription rather than on/off switching.

### 3.6 H3K4me3 is the most important predictor of gene expression

Integrating all the data generated, we started by characterizing their distribution over all genes categorized by two factors, namely gene expression and gene length. Figure 7A shows the input normalized profiles of different epigenetic marks, and GC content based on the gene expression groups. Genes in heatmaps are sorted by gene length. MNase, Pol II, H3K4me3 and H3K9ac show accumulation in the 5′-CDS in expressed genes with decreasing intensity along the ORF. However, most signals are still high and correlate to gene expression level in the 3′-CDS. The 5′-accumulation is not that pronounced in H3K27me3, which shows more equal distribution along the ORF. Hence, we further investigated how the epigenetic marks are distributed along the gene structure, based on their length. MNase signals show a strongly phased pattern in all categories of gene expression, which is evidently seen when the genes are sorted by length. Supp. Fig. 10A shows a strong positive correlation of exon length and nucleosome counts in exons. Similarly nucleosome occupancy is positively correlated with gene expression (Fig. 7A). Similar to the strongly phased signals of MNase, we observe that Pol II signals are also phased and show positive association with gene expression.

**Figure 7.**
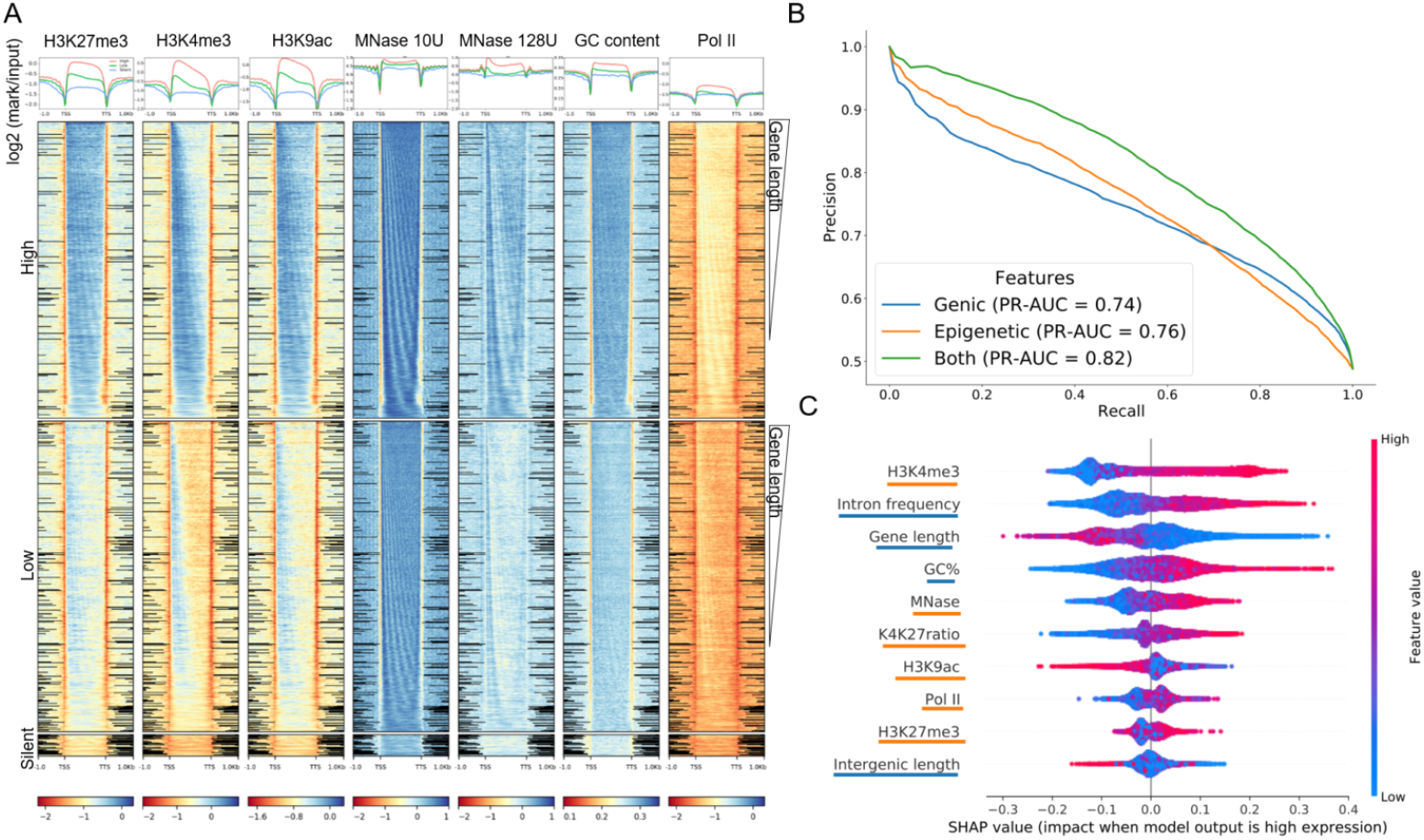
Prediction of gene expression by epigenetic marks. A, Distribution of epigenetic marks in different transcriptomic groups. The input normalized enrichment profiles (y-axis; top) of different epigenetic marks are shown. The enrichment values are shown as a heatmap (below). Genes (rows) are split into three categories based on gene expression: High (TPM>2), Low(0<TPM<2) and Silent (TPM=0), and are sorted by decreasing order of gene length in each. Distance shown on the x-axis is scaled, i.e. all genes (TSS-TTS) are either stretched or shrunken to a length of 1500 bp, adding 1000 bp up- and downstream of the gene. Enrichment profiles and heatmaps were plotted using deeptools2. B, Results of classifying Low and High gene groups using different data (features Genic: related to gene structure, Epigenetic: using abundance of histone marks and MNase, Both: Genic and Epigenetic). Precision-Recall curve with average values from a 40-fold cross validation with random forests indicating features by different colors. C, Analysis of feature importance using both Genic and Epigenetic features (underlining color indicates type on y-axis, see legend in B). Features are listed in decreasing order of classification importance from top to bottom. The importance (SHAP value, x-axis) of a feature for each gene illustrates its contribution to classification as High or Low, with positive and negative SHAP values, respectively. The colour gradient depicts the feature value in scale from low to high, e.g. the length of a gene (third row). For example, long genes strongly contribute to the prediction of lowly expressed genes. The overlapping dots are jittered in the y-axis direction.

Interestingly, all epigenetic marks are consistently low at 5′-and 3′-non coding regions showing a clear gap in all analyses, thus fostering the assumption that intergenic regions hardly contribute to gene regulation. All silent genes have very faint signal of all epigenetic marks, supporting our conclusion that lowly occupied nearly naked DNA is a hallmark of gene inactivation in *Paramecium*.

The visualization in the heatmaps in Fig.7A reveals a phasing pattern for almost all marks, as genes are ordered by gene length in each expression group. This means that nucleosomes are indeed well positioned in all ORFs and along the entire length, but with varying intensity, due to differences in gene expression. As one will have assumed then that the histone marks need to follow the nucleosome pattern, this follows also the GC content oscillations in position and quantity. As such, this cis-factor may be involved in predetermining nucleosome positioning and consequently gene expression. We investigated effects of gene length and mRNA levels, and observed that shorter genes show higher mRNA levels (Suppl. Fig.10B), and as such gene length itself appears to be a factor limiting transcriptional efficiency. Surprisingly, we observe the phasing pattern also for Pol II occupancy. This would suggest that Pol II shows association with nucleosomes along the entire ORF, and interestingly the higher Pol II occupancy in highly expressed genes does not indicate that this association is a mechanism of transcriptional inhibition. In agreement with the conclusion from the pausing index analyses, this Pol II nucleosome association appears to be a mark of highly expressed genes, although one could get the impression that Pol II stops at every single nucleosome, which could also be an argument for inefficient elongation.

As we observed some intriguing patterns of histone marks, especially of H3K27me3 which is abundant in highly expressed genes, we checked the correlation of all epigenetic marks with each other with mRNA (Suppl. Fig. 11A). We observed that all epigenetic marks are positively correlated (Pearson correlation > 0.6) with each other, and a bit weaker with mRNA (Pearson correlation > 0.30). We wondered what the individual contribution of gene structural characteristics and occupancy of epigenomic marks is with respect to gene expression. Thus, we constructed a machine learning classifier to predict genes as highly or lowly expressed using epigenetic features and genic features (see Methods). After experimenting with different classification methods (data not shown), our final model is based on a random forests algorithm, which accurately predicts gene expression with an average PR-AUC of 0.74 and 0.76 for genic or epigenetic features, respectively. The model combining all information performed best (PR-AUC of 0.82, Fig.7B). These differences where statistically significant (Suppl. Fig.11B). Experiments in Fig.7B where done using histone marks in the complete gene body. When quantification is restricted to the proximal TSS region (TSS+300 bp), performance decreased (Suppl. Fig.11C), supporting a role of those marks through out the gene body.

Further, we interrogated the best performing model on the importance of each feature in obtaining the classification (Fig.7C). According to the feature importance values calculated on our best performing model, H3K4me3, intron frequency, and gene length are the top three features required to classify gene expression. Intergenic length, and H3K27me3 are among the least important features for our model. The presence of H3K27me3 in the whole gene body; its high correlation to other histone marks, and highly expressed genes does raise the question of the role of H3K27me3 in Mac chromosomes of *Paramecium*.

### 3.7 H3K4me3 and H3K27me3 co-occur at plastic genes

We consequently asked for the contribution of individual features to gene regulation. We utilized RNA-seq data from environmental states that include four different serotypes at different temperatures, starvation, heat shock, and cultivation at 4°C [12]. Using those data we dissected genes showing large expression variations (high plasticity) during vegetative growth in different environments to identify dynamically regulated genes from housekeeping genes (see Methods and Suppl. Fig.12). We defined four classes of plasticity (G1-G4), where G4 genes showed the largest amount of variation. We again used the random forest algorithm to analyze whether genic/epigenetic factors contribute to the accuracy of gene expression prediction for each gene plasticity group. The performance of expression prediction decreased for genes with higher plasticity (Fig.8A). Thus, plasticity of gene expression seems to be accompanied with additional and unknown features contributing to gene regulation.

**Figure 8.**
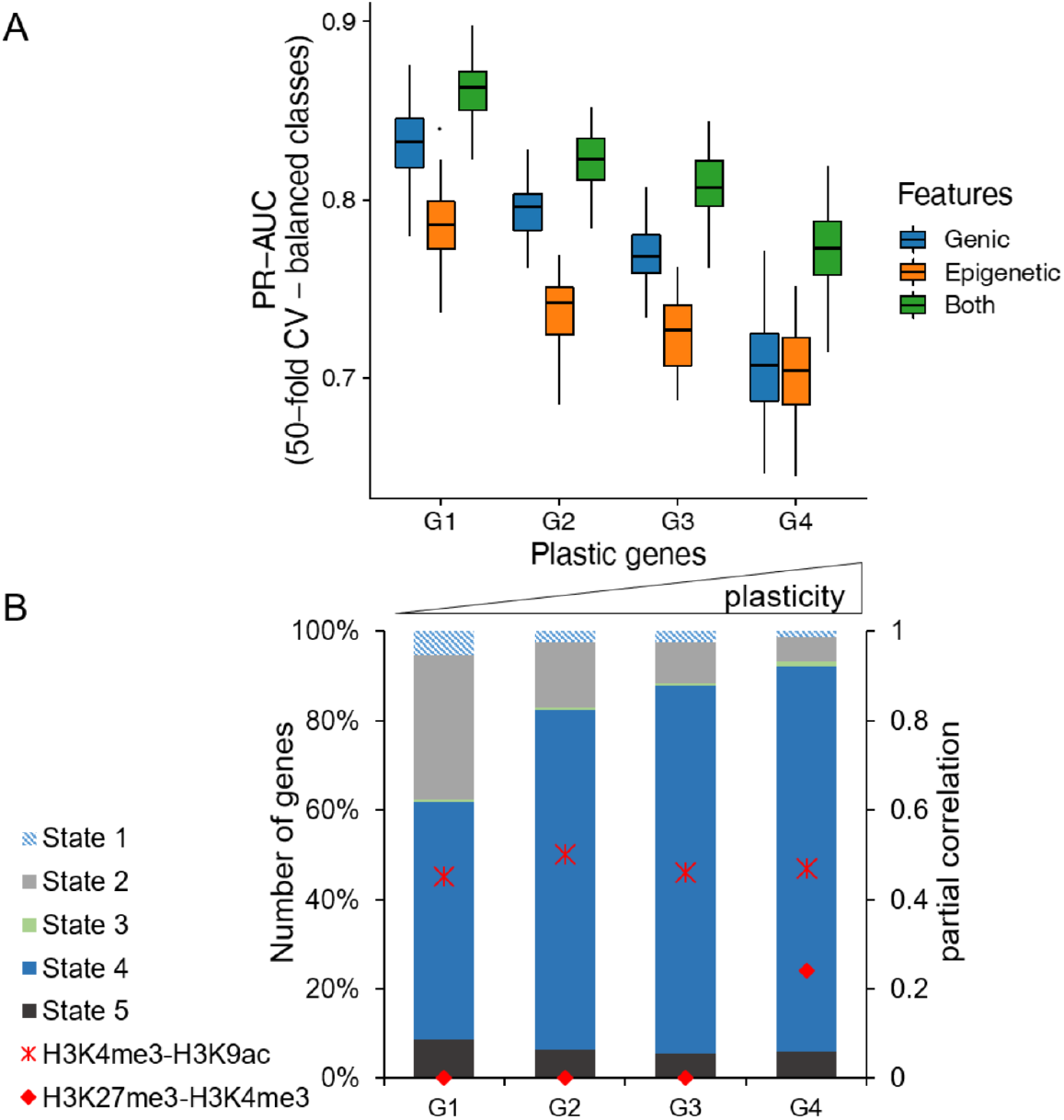
Prediction of gene expression for genes with high plasticity. Genes were separated into four groups by their plasticity, which is defined by a large variation in gene expression among different conditions. A, Box plot showing the distribtion of classifier performance values for genes with different plasticity (50-fold CV based PR-AUC) for the same three feature sets as in Fig.7B. The number of genes in each plastic gene group was randomly subsampled to have equal number of genes in high and low expressed category. B, Distribution of chromatin states among plastic gene groups. We only included genes with a ChromHMM state overlap of at least 80% (see Fig. 5). Additionally, partial correlation values for H3K4me3-H3K9ac (cross) and H3K4me3-H3K27me3 (circle) are shown in red for each group.

TO get further insights, we checked the chromatin states based on our ChromHMM segmentation of the four categories of plastic genes (Fig.8B). These show gradual differences with most apparent increase of CS4 and decrease of CS2. This suggests, that epigenetic marks are not only used for control of gene expression but moreover for gene regulation. We studied the differences of histone marks of these categories in more detail and calculated the partial correlation between different modifications (see Methods). Fig.8B shows an increase in partial correlation of H3K4me3/H3K27me3 for the most plastic genes only, suggesting that the interplay between histone marks varies in the four considered groups.

## 4 DISCUSSION

### 4.1 Genomic and epigenomic paradoxes

At first glance, the genomic structure of the *Paramecium* Mac seems paradox. Although *Paramecium* is extremely gene-rich, with approx. 40.000 genes [5], the size limitations of intergenic regions and introns provide only restricted capacity for differential gene regulation. This is different compared to genomic/epigenomic features in metazoens, because unicellular organisms do not need to differentiate into distinct tissues with all the known epigenetic manifestations to guarantee for cell type specific gene expression patterns. However, the *Paramecium* epigenome still needs to manage dynamic regulation of gene expression and proper transcription of mRNA. We know that histone marks do not just control condensation and transcriptional on/off switches, but interact with capping enzymes, splicing factors and elongation factors to guarantee for mature mRNA synthesis [33].

Thus, we aimed to answer the question in which manner the Mac epigenome signature is associated with transcriptional regulation in this ciliate. In *Paramecium*, nucleosome occupancy, and as a result histone modifications, appear to be associated in general with active transcription, because segmentation of MNase and ChIP data shows a large number of genes where our setup detects only low or no signals (CS4 in Fig.5). Correlation of this chromatin state with gene expression, indicates surprisingly, that low nucleosome occupancy, regardless of the histone marks, is associated with silent or lowly expressed genes. One could therefore interpret naked or lowly occupied DNA as a default state, which needs to be occupied with nucleosomes first to become transcribed into mRNA. As such, the epigenome of *Paramecium* appears paradox as well, as gene inactivation becomes realized by low nucleosome occupancy and this is contrary to the classical models.

Textbooks describe gene inactivation by a hierarchical chromatin folding from open 10nm fibres to condensed and higher occupied 30nm filaments. Active transcription accompanied by open, accessible chromatin in mammals was highly supported in the last years by many studies of DNA accessibility using ATAC, NOMe, DNAse-Seq or methods free of enzymatic steps like sedimentation velocity centrifugation [30,34,45]. Our data does not support this model for *Paramecium* Mac chromatin suggesting a totally different chromatin associated mechanism of gene inactivation. This seems surprising and raises many more questions how in particular spurious and aberrant transcription of PolII in open regions is inhibited or whether this could be tolerated to some extent.

In most species, condensation of chromatin is accompanied with linker histone H1 recruitment and studies on *Drosophila* chromatin demonstrate H1 occurring exclusively at closed heterochromatic loci [44]. We are not able to identify a macronuclear histone H1 variant in *Paramecium* supporting the idea of condensation-free gene inactivation. To be precise, we have to distinguish macronuclear and micronuclear linker histones in ciliates. *Tetrahymena* has distinct Mac and Mic specific H1 histones, where the macronuclear version (Hho1) is non-essential [54]. Hho1 knockouts show an overall decondensation of macronuclear chromatin [29]. As such, the lack of *Paramecium* Hho1 homologs fits to our finding and moreover suggests differences in the chromatin organization between *Paramecium* and *Tetrahymena*.

### 4.2 Bistable H3K4/K27me3 as a mark of poised genes?

Another question we followed is whether the H3K27me3 could be involved in gene inactivation. Our ChIP data does not suggest H3K27me3 to be associated exclusively with silent or lowly expressed genes. Asking for the function of this modification in the vegetative Mac, its role is unlikely the condensation of chromatin and the segmentation shows H3K27me3 co-occurring in varying ratios with the H3K9ac and H3K4me3. Our data suggests that genes with high regulation dynamics show an increasing correlation for H3K27me3 and H3K4me3. This is one of the best studied bivalent domains for poised chromatin where chromatin is placed into a waiting state for future activation and this was described to occur in particular in embryonic stem cells [46, 66]. There is an ongoing debate whether poised chromatin is bi-stable or bivalent, the latter representing a background population of fragments with active and silent marks, whereas bi-stability means the frequent switching between monostable active and silent states [58].

The polyploidy of the *Paramecium* Mac introduces here an additional layer of complexity. Similar to ChIPs of different cell states from a culture of metazoen cell cultures, which cannot dissect different cell states of a mixture from a real bivalent domain, we cannot be sure here, that the 800 copies of a gene in the Mac are co-regulated. If *Paramecium* for instance would use gene dosage to regulate gene expression level, one would expect different ratios of marks: some copies silent, some copies active. This is what we can observe to some extent, because the random forest analysis suggests the K4/K27me3 ratio to explain the gene expression level better than the H3K27me3 alone. In a previous study, increased H3K27me3 levels in association with decreased levels of H3K4me3 at an endogenous reporter gene have been shown to go along with siRNA mediated silencing [25], which supports the K4/K27me3 ratio hypothesis for controlling gene expression levels. In addition, the finding, that we see increasing partial correlation values of K4/K27me3 in genes which show high regulation dynamics could be called poised as such. This suggests that the bivalency of K4/K27me3 in chromatin poising could be an ancient and general mechanism, rather than an invention of metazoens.

In *Paramecium* the polycomb group methyltransferease Ezl1 was demonstrated to mediate both H3K9me3 and H3K27me3 during development: loss of these marks are accompanied by loss of transposon repression and elimination and in addition a transcriptional up-regulation of early developmental genes [22]. As Ezl1 shows also low expression during vegetative growth, it remains to be elaborated whether Ezl1 or another SET-domain containing enzyme catalyzes the replicative maintenance of H3K27me3 during vegetative cell divisions.

From an evolutionary point of view this could imply that although *Paramecium* is unicellular, the epigenomic repertoire already has the capacity to manifest vegetative gene expression regulation during development, meaning to place histone marks for poising genes. Inheritance of gene expression pattern was previously shown also for the multigene family of surface antigen genes as transcription of a single gene follows the expression pattern of its cytoplasmic parent [6, 55] but we would need to analyse the genome wide extent of such an inheritance, and/or whether such a mechanism is coupled with other genomic parameters, like for instance sub-telomeric localization of the respective genes.

### 4.3 ChIPseq reveals broad domains instead of narrow peaks

Looking for the distribution of marks along genes, the absence of narrow peaks becomes apparent as all histone mark distributions are more comparable to broad domains instead of local and narrow peaks, which explains the failure of peak calling. Broad domains were also found in higher eukaryotes. For instance, H3K27me3 was shown in mammalian chromatin to be distributed along ORFs [67]. Also in mammals, broad H3K4me3 was demonstrated for tumor-suppressor genes with exceptionally high expression, where this mark has also been attributed to transcriptional elongation [13]. In addition to tumor-suppressors, broad H3K4me3 domains have been implicated with genes for cellular identity and transcriptional consistency; as the broadest domains show increased Pol II pausing, the authors suggest the broad mark as a buffer domain to ensure the robustness of the transcriptional output [8].

This model could also fit to our observations, which do not only suggest H3K4me3 as the key regulator of transcription, but that H3K4me3 appears in broad domains along ORFs highly covered with Pol II. With respect to the different patterns of Pol II along ORFs compared to other species, either for poised or non-poised genes, the buffer domain model could hold true for the majority of *Paramecium* genes.

### 4.4 Nucleosome positioning and GC content

*Paramecium* has an exceptional genome composition with an average GC content of 28% and including the even more AT-rich intergenic regions. It is known that GC content favors nucleosome positioning [61]. Our data shows that nucleosome occupancy is mostly restricted to ORFs, which would correlate to increased GC-levels, but also correlated to gene expression levels as higher expressed genes show higher occupancy of promoter proximal- and intron-associated nucleosomes. It is difficult to reason in how far the sequence content in the *Paramecium* genome itself encodes the deposition of nucleosomes from our data. There is ample discussion about the DNA sequence preferences of nucleosomes [43] and also MNase-seq can generate a signature of higher occupancy at GC-rich regions, on naked as well as occupied DNA [16]. One may conclude, that this bias explains the large drop of MNase-seq read occupancy at intergenic regions. However, analysis of ChIP-seq data show a similar drop at intergenic regions and similar phasing patterns in our data and Suppl. Fig.13 suggests that our procedure and the applied PCR amplification have minimized GC biases. We argue that it is unlikely to observe these trends exclusively due to methodological biases in AT-content.

Our results of nucleosome positioning fit to observations in *Tetrahymena* where well-positioned nucleosomes in the Mac match GC-oscillations, but are also affected by trans-factors, e.g,. the transcriptional landscape [63]. In addition, studies in *Tetrahymena* revealed that N6 methyladenine (6mA) is preferentially found at the AT-rich linker DNA of well positioned nucleosomes of Pol II transcribed genes [40, 62]. Also in *Paramecium* 6mA sites enriched between well positioned nucleosomes are positively correlated with gene expression [28]. The latter finding would fit to our observations: the more nucleosomes, the more 6mA, the more transcription.

### 4.5 Qualitative aspects of gene expression

In order to understand the relation between epigenomic data and gene expression, throughout this study we categorized genes based on their expression levels (high, low, silent). While this categorization helps, it should be treated with a grain of salt as the cut-offs are rather arbitrary. Another aspect which requires cautious interpretation is the analyses presented in Figure 7. Specifically, Figure 7A shows the linear relation between epigenetic signals and mRNA expression in a qualitative manner. The random forests analysis, presented in Figures 7B and C, reveals both the linear and non-linear relationships inherent in the epigenetic data while calculating the probabilities to predict/classify a gene as highly or lowly expressed. For example, we can observe that H3K9ac is directly proportional to the different expression groups in Figure 7A. However, Figure 7C suggests genes with low H3K9ac to be associated with high expression. While this may seem counter intuitive, both results are correct owing to the high collinearity of epigenetic marks (Suppl. Fig.10A). Hence, the random forests model relies on the H3K9ac signal only when the H3K4me3 signal is not sufficient to increase the probability of predicting a gene as highly expressed.

### 4.6 A divergent mechanism of transcriptional elongation

How can the highly regulated CTD phosphorylation and interaction with the different RNA modification and elongation complexes of higher metazoens be compared to our data? *Paramecium* Pol II does not exhibit the serine rich heptamer repeats. Thus, it would be surprising if a regulated and patterned phosphorylation of individual serines would be possible. As the Paramecium CTD is still rich in serines, although not organized in a heptamer repeat structure, it still seems likely that phosphorylation could be an activating mark. It seems quite tempting to speculate that Pol II of *Paramecium* does not need to be that highly regulated compared to mammals. First of all, alternative splicing is extremely limited and no single example of exon skipping has been reported [32], and therefore the well positioned nucleosomes do not need to control this. As intron nucleosomes could still be involved in splicing efficiency this may also not be necessary as *Paramecium* introns are apparently recognized by intron definition and even artificially introduced introns in GFP are efficiently spliced [32]. Our data can be interpreted in that introns serve as a nucleosome positioning place maybe to attract more introns to the gene thus supporting transcription. This would be supported by our data showing that genes with higher intron frequency show higher transcript levels.

Concerning the issues of pausing and elongation, our data suggests pausing to occur, but the pattern is different to other species because we find high levels of Pol II associated with nucleosomes along the entire ORF not only restricted to +1 nucleosomes. Given the fact that +1 nucleosomes are quite prominent, the question raises whether the stops of Pol II at +1 nucleosomes are mechanistically different from stops at all nucleosomes inside the ORF, or whether this is a general phenomenon of *Paramecium* Pol II to stop at nucleosomes, maybe by less efficient elongation. For instance, the tiny introns of *Paramecium* do not contribute to a significant enlargement of transcriptional units compared to other species with introns which are often much larger than the exons. It is therefore the question whether Pol II elongation has the need to be highly supported. In fact, *Paramecium* and *Tetrahymena* miss homologs of NELF, and two recent studies demonstrated the mediator complex, a key regulator of Pol II interaction with transcription and elongation factors, to be highly divergent in *Tetrahymena* [24, 60]. *Addionally, in Paramecium* we cannot identify all components of the Paf complex regulating elongation, 3′-end processing and histone modification in lower and higher eukaryotes [31]. Especially, the subunit Paf1, involved in serine phosphorylation of the CTD of Pol II, is missing and which fits to the missing serine repeats of the CTD. Because of the lack of canonical elongation systems going along with a lack of conserved serine residues, we conclude that transcriptional elongation in *Paramecium*, is regulated in a different manner. As a result of this, we can observe the high PolII occupancy in highly expressed genes in our data, which we would not expect in metazoens. As discussed above, broad H3K4me3 going along with increased occupancy of Pol II in ORFs might be an alternative control of transcription by buffer domains. It seems tempting to speculate this strange form of Pol II buffering represents an alternative or maybe an ancient form of elongation control.

## 5 CONCLUSION

This is the first description of the *Paramecium* vegetative chromatin landscape which appears to be quite different to that of higher and other unicellular eukaryotes. Broad domains along the gene bodies apparently regulate transcription whereas the non-coding and non-expressed regions are devoid of epigenetic information. Paradoxically, our data also indicates silent genes to be devoid of epigenetic information and it has to be clarified if and how the cell prevents spurious Pol II activity at these unoccupied regions. The Pol II distribution we observe is also quite different to other species, the process of transcriptional initiation and elongation appears to be controlled without sophisticated control of CTD phosphorylation and canonical complexes, like NELF, Paf, and Mediator that assist Pol II in generating mature mRNA. However, this work here attributes to the vegetative nucleus, only. We have to keep in mind that the transcriptional machinery needs to switch its mode of action to lncRNA transcription from the meiotic micronuclei during development. As such, functional and temporal dynamics require more alterations of the polymerase complex than in other species. There are plenty of challenges left, especially about the control of PolII without or with limited CTD phosphorylation. Our study shows the unusual pattern of Pol II in expressed genes and in the light of so many missing interaction partners of Pol II, its not a surprise that the epigenome looks different to other species in addition to the fact that no mitotic condensation is necessary in the Mac. Concerning Pol II interaction complexes, future studies will need to show whether some components are absent, or whether they are too divergent such that reverse genetics cannot identify them. Their identification and contribution to PolII activity and modulation will shed light into the mechanisms controlling mRNA and lncRNA transcription and the epigenetic marks in support of them.

## Acknowledgments

This work was supported by grants from the German research Council (DFG) to MS (SI1379/3-1), MHS SCHU (3140/1-1) and MJu MJ (CRC894). AS was supported by the German Federal Ministry of Research and Education grant for de.NBI (031L0101D). We are grateful to Laura Arrigoni and Ulrike Boenisch for support with NEXSON and ChIP-seq. We are grateful to Sandra Duharcourt and Melody Matelot for sharing unpublished MNase datasets. The authors acknowledge the support of the Freiburg Galaxy Team, University of Freiburg (Germany) funded by the Collaborative Research Centre 992 Medical Epigenetics (DFG grant SFB 992/1 2012) and the German Federal Ministry of Education and Research BMBF grant 031 A538A de.NBI-RBC.

## Notes

### Competing Interest Statement

The authors have declared no competing interest.

