## Supplementary Figures for "Broad domains of histone marks in the highly compact *Paramecium* macronuclear genome"

**Suppl. Fig. 1**

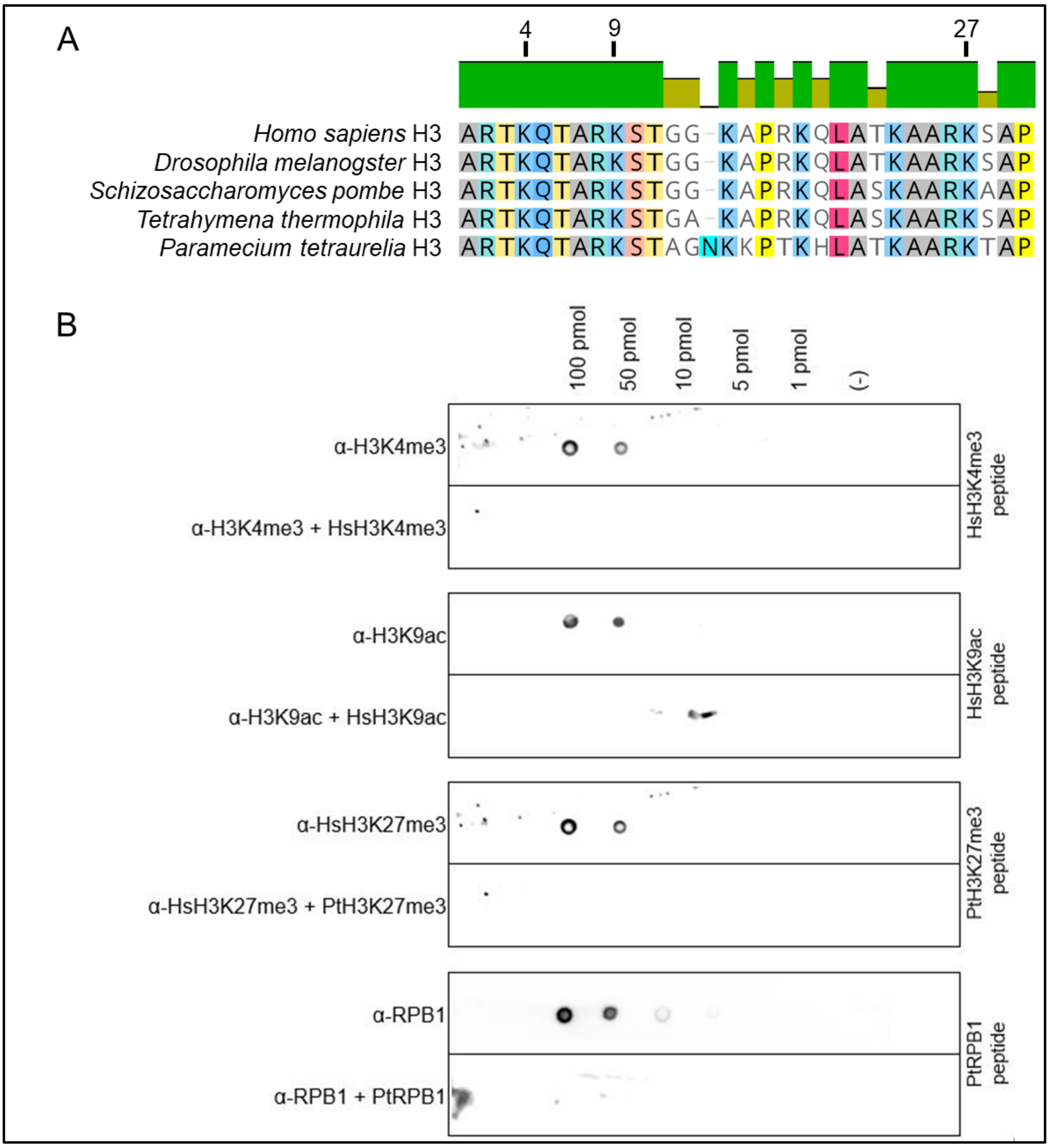


**Suppl. Fig. 1** A) Sequence alignment of the N-terminal amino acid residues (1-30) of histone H3 proteins of indicated species. Positions analyzed for their modifications are highlighted (H3K4, H3K9 and H3K27). Accessions: CAB02546 (*Homo sapiens*), NP_001027285 (*Drosophila melanogaster*), P09988 (*Schizosaccharomyces pombe*), XP_001016594 (*Tetrahymena thermophila*) and PTET.51.1.P1080178, H3P1 (*Paramecium tetraurelia*). B) Competition assay. 100 pmol to 1 pmol of each peptide were spotted and either probed with the corresponding antibody or with antibody that was blocked with the indicated peptide in a 10 fold excess in advance. For all antibodies, blocking results in a loss of specific binding to the membrane bound peptide. Blocking the α-HsH3K27me3 with the Paramecium PtH3K27me3 peptide also results in complete competition.

### Suppl. Fig. 2

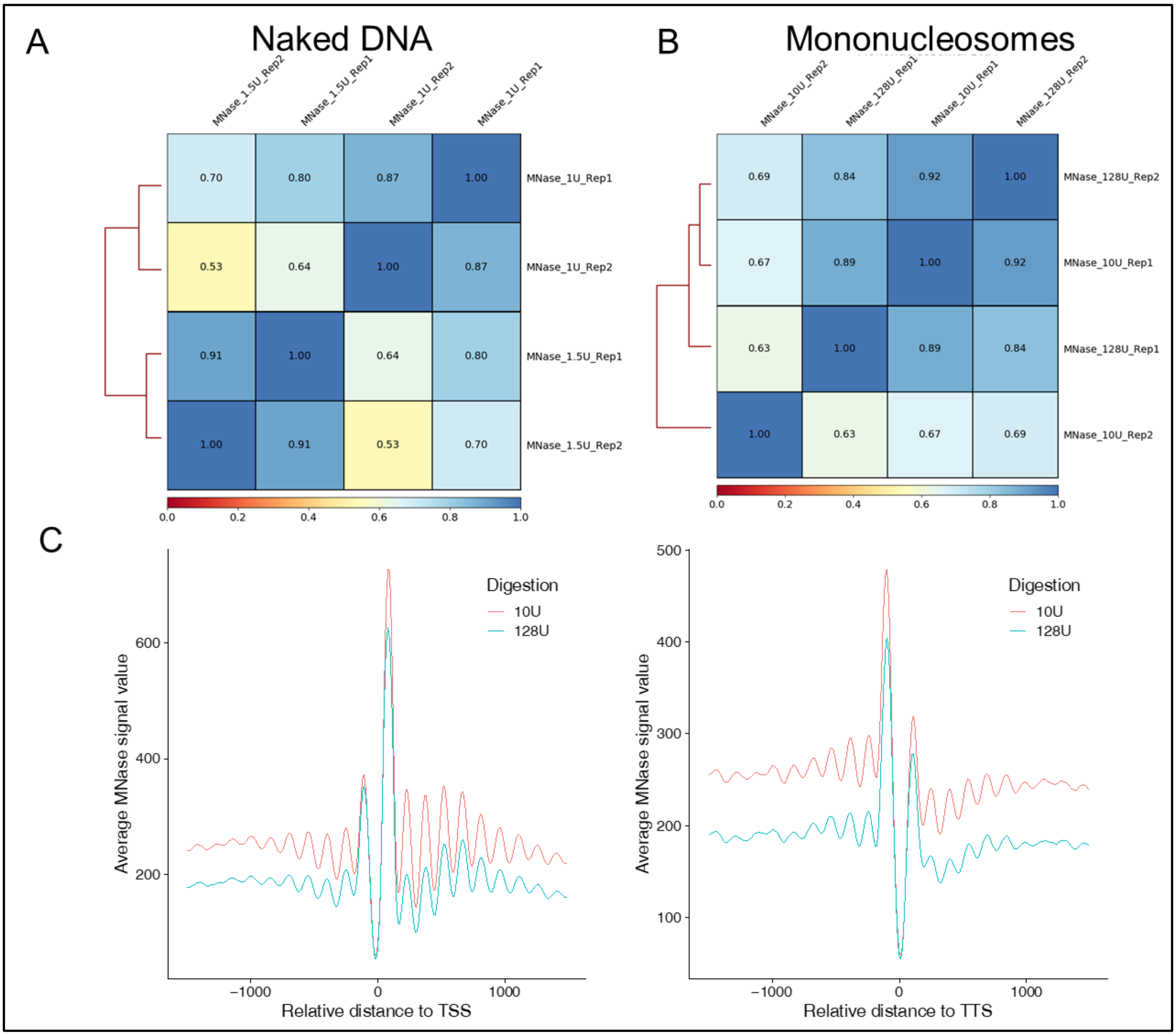


**Suppl. Fig. 2** Pearson correlation between naked DNA (A) and MNase-seq (B) replicates of mononucleosomes obtained by different digestion units. C) Profile plot of nucleosome distribution at the transcription start site (TSS), and the transcription termination site (TTS) respectively.

### Suppl. Fig. 3

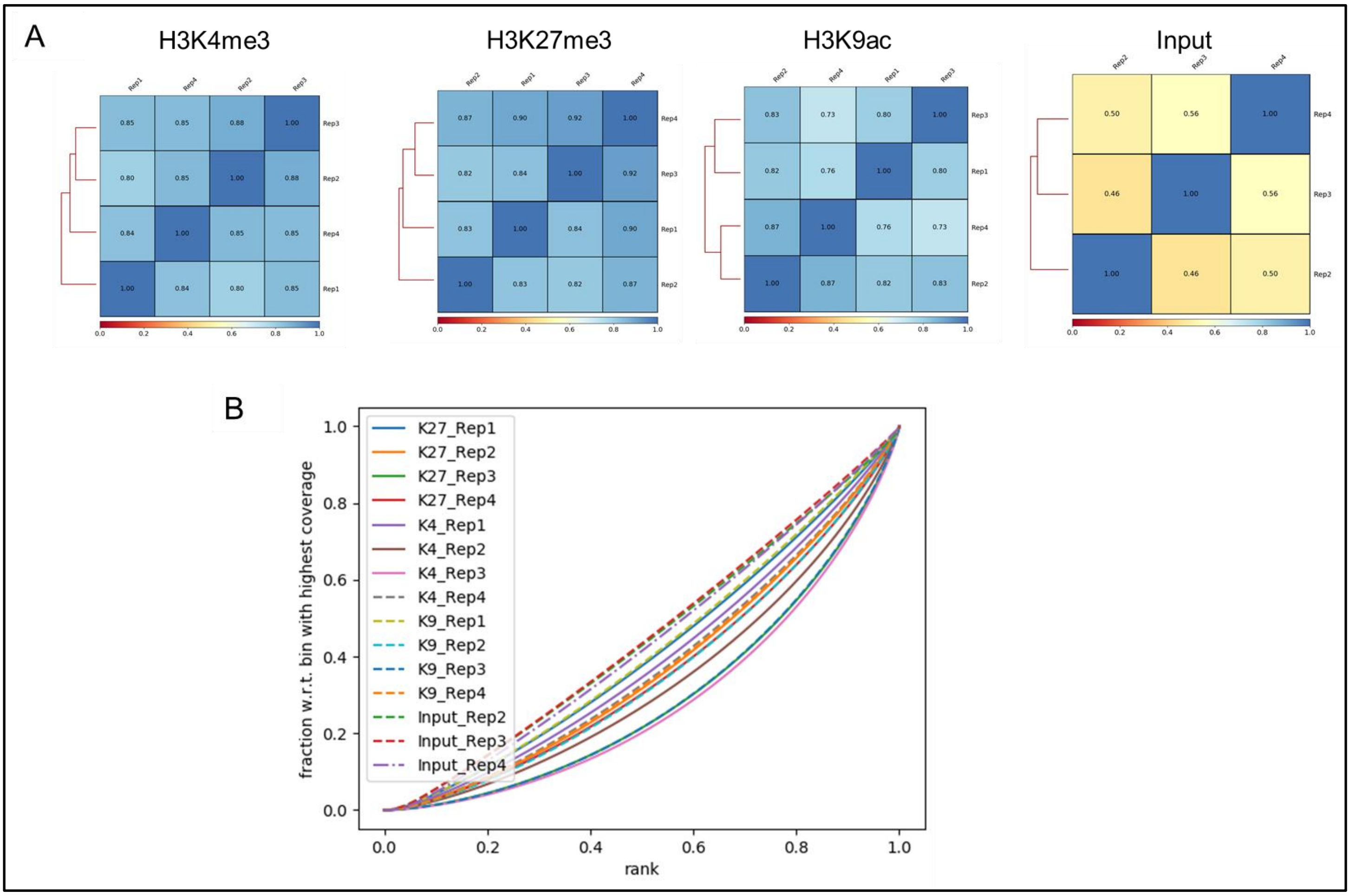


**Suppl. Fig. 3** A) Subsampled Pearson correlation of read counts between histone mark and input replicates. B) Fingerprint plot, generated with deeptools, showing the quality of replicates.

### Suppl. Fig. 4

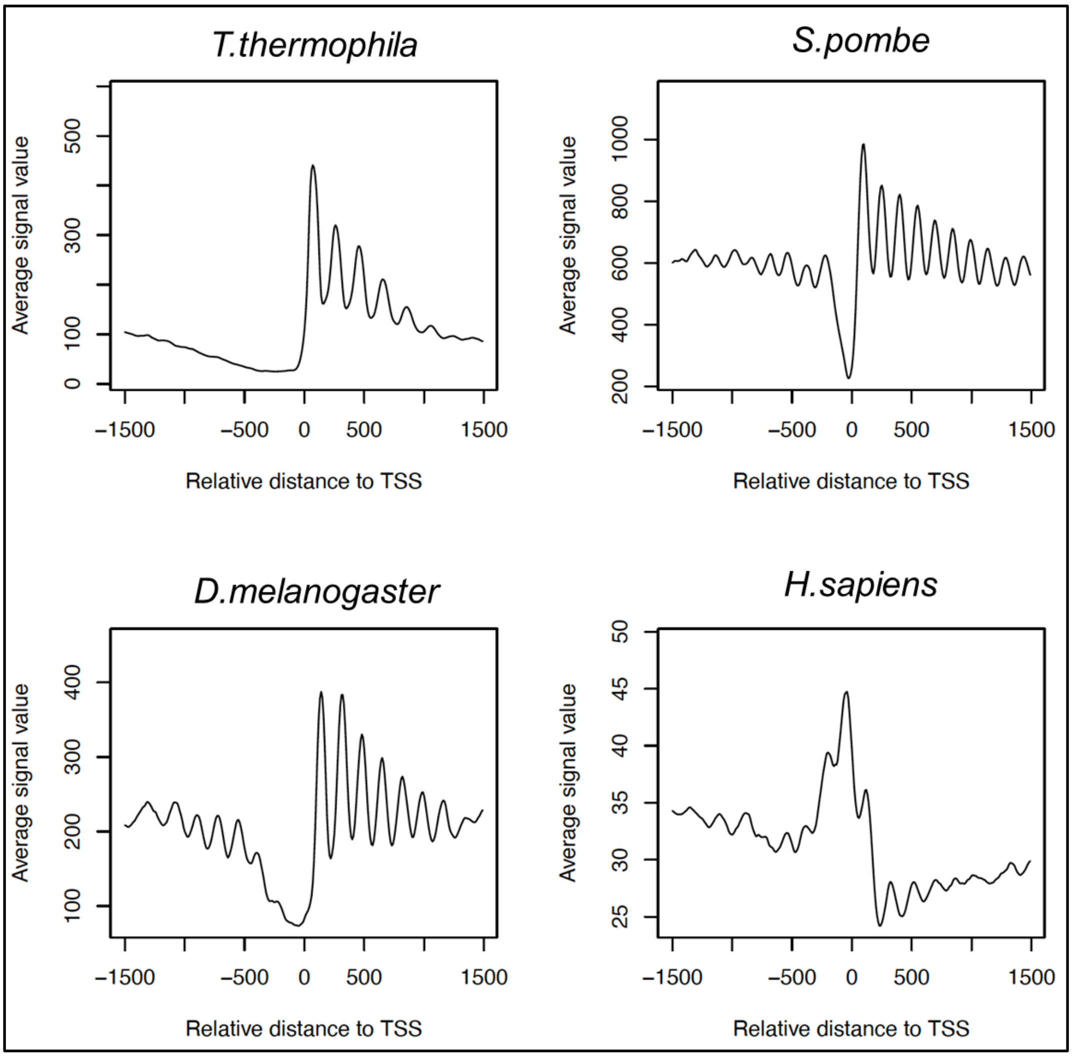


**Suppl. Fig. 4** Profile plot of nucleosome distribution at the transcription start site (TSS) for all analyzed genes of *Tetrahymena thermophila*, *Schizosaccharmoyces pombe*, *Drosophila melanogaster* and *Homo sapiens*. Signals for 1500 bp up- and downstream of the TSS are shown. Data set information is found in Supplementary Table 1 (Comparative MNase analysis).

### Suppl. Fig. 5

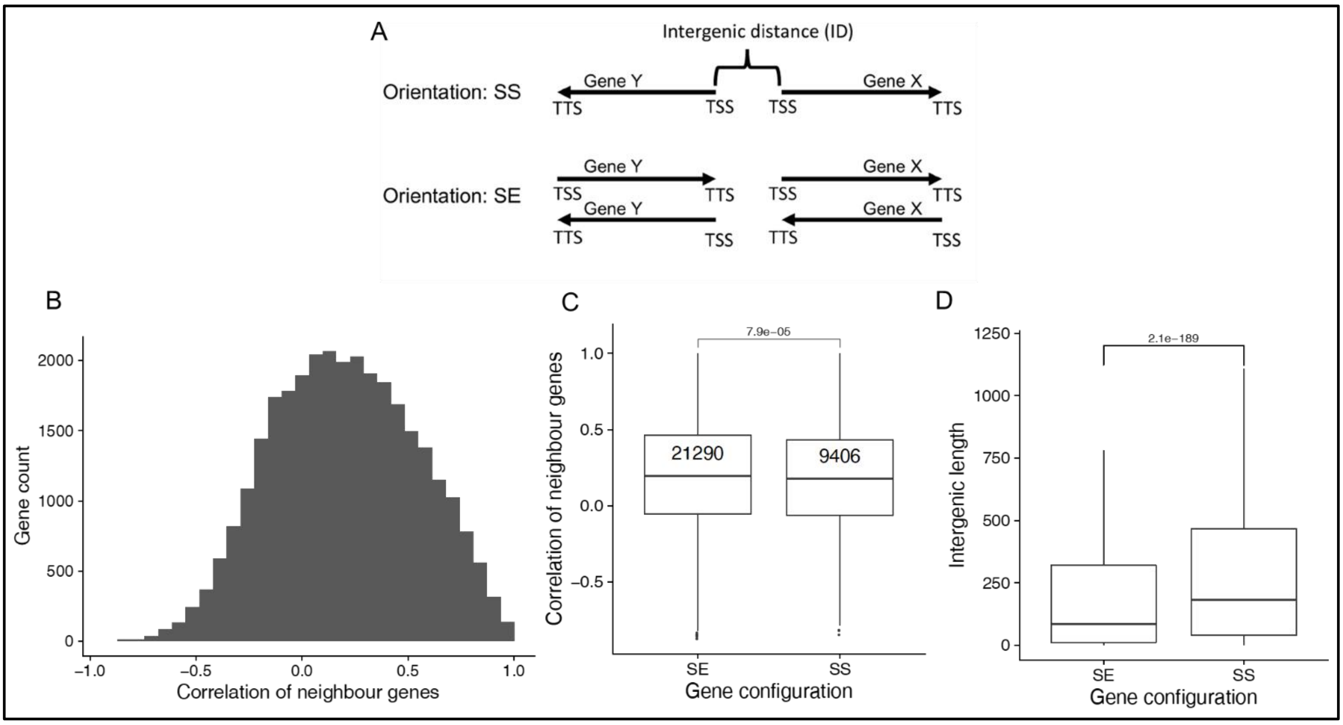


**Suppl. Fig. 5** A) An illustration of the gene orientation-based grouping. SS = bidirectional, SE = unidirectional B) Distribution of the Pearson correlation of neighbouring genes expression from different serotypes. Data reused from our publication (Cheaib et al, DNA Res 2015). C) Pearson correlation of neighbouring genes expression for genes with different configurations. D) Length of intergenic regions is shown for genes with the same configurations as in B. Y-axis is zoomed in and outliers are not shown. The P-values shown are based on a two-tailed Wilcoxon test.

### Suppl. Fig. 6

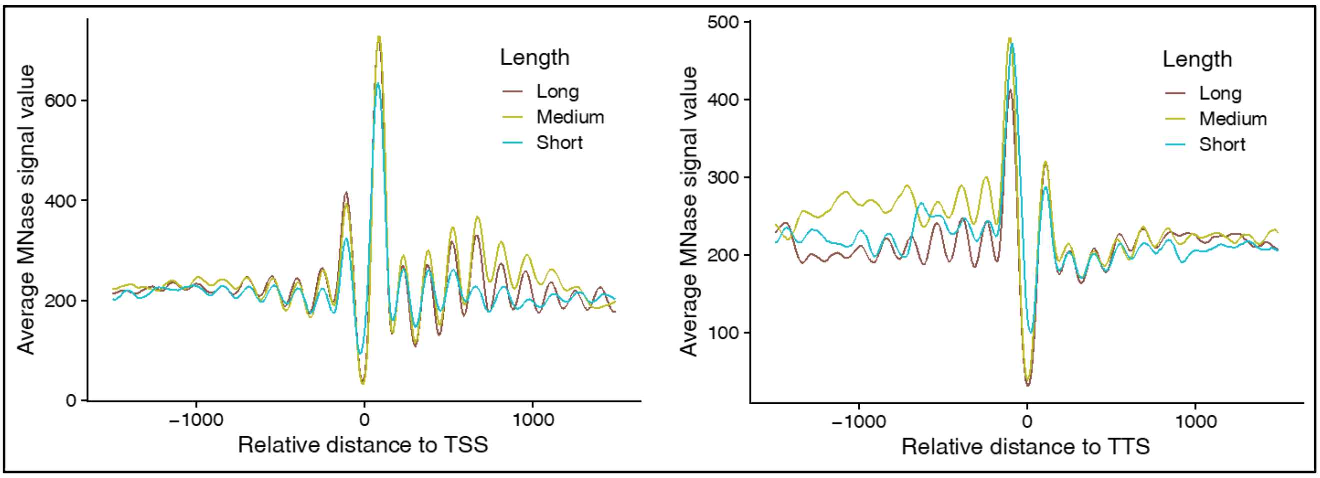


**Suppl. Fig. 6** Profile plot of the nucleosome distribution at the TSS and TTS, stratified according to gene length groups (Short: 50-765 bp; Medium: 765-1470, Long: >1470).

### Suppl. Fig. 7

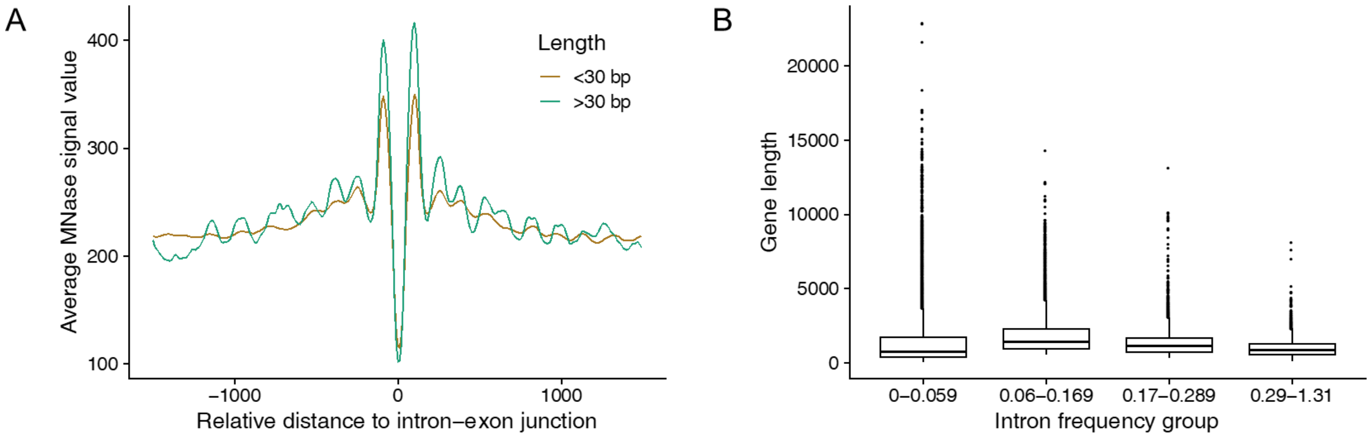


**Suppl. Fig. 7** A) Profile plots of nucleosome distribution around the intron-exon junctions, categorized based on intron length groups. B) Box plot showing the distribution of gene length across different intron frequency groups (as shown in Fig.4). A Krustal-Wallis test showed that the expression distribution of all intron frequency groups is statistically significant ($P < 2.2e-16$).

**Suppl. Fig. 8**

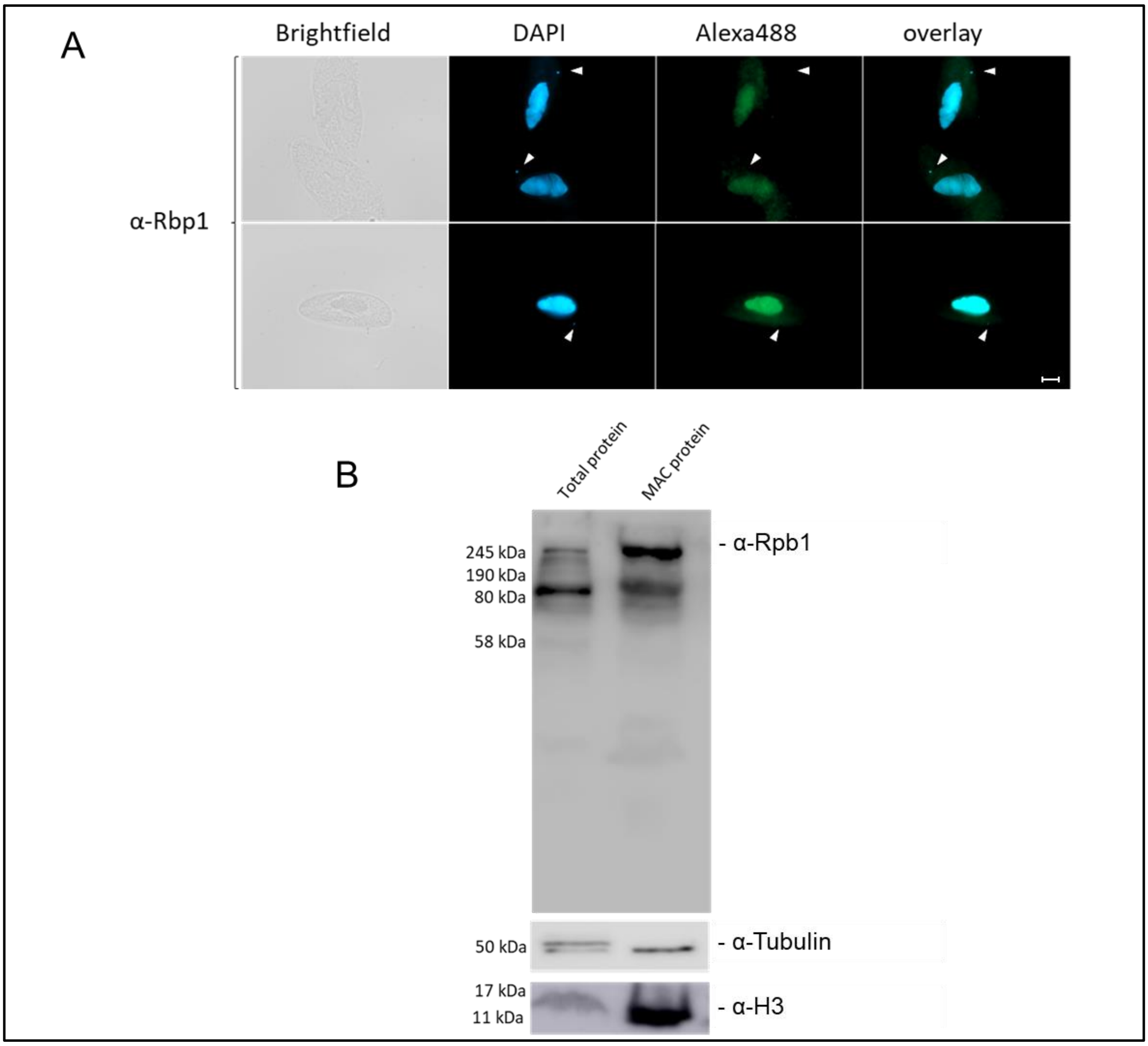


**Suppl. Fig. 8** Localization of Polymerase II by immunofluorescent staining and western blots. A) Vegetative Paramecium cells were stained by indirect immunofluorescent staining using custom antibody directed against Rpb1 as the largest subunit of Polymerase II. Primary antibody was labeled with Alexa488-conjugated secondary antibody (green) and nuclei were stained with DAPI (blue). Arrowheads point at micronuclei. Representative overlays of Z-Stacks are shown. Scale bar is 10 µm. B) Western blots using custom antibody against Rpb1 were performed as previously described (Klöppel et al., 2009). Protein lysate from whole cells (total protein) and protein from fractions of enriched macronuclei (MAC protein) were blotted and the membrane was decorated with antibodies against Rpb1 (200 kDa), α-Tubulin (50 kDa) and Histone H3 (15 kDa) as loading control.

### Suppl. Fig. 9

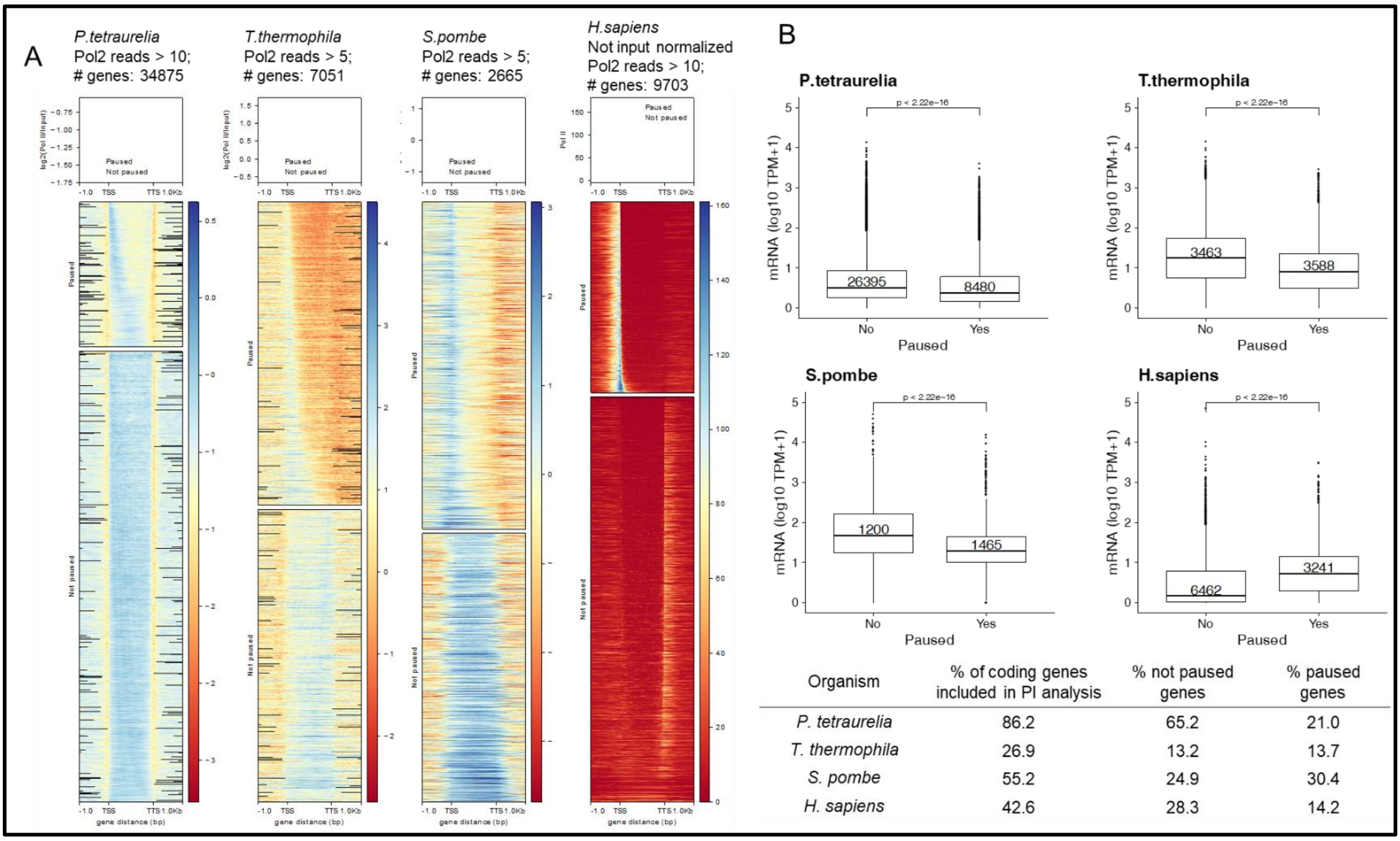


**Suppl. Fig. 9** A) A heat map and profile of the Pol II enzyme in different organisms (see Supplementary Table 1: Comparative Pol II analysis sheet). The number of genes used in the plots are shown above each plot, after applying filters on the number of reads to avoid bias from silent genes. The genes are categorized as paused, if they had a pausing index more then 1.5 (see Methods). B) A box plot of the gene expression stratified according to their paused status in different organisms is shown. Table below shows, how many genes are included in the pausing index analysis.

### Suppl. Fig. 10

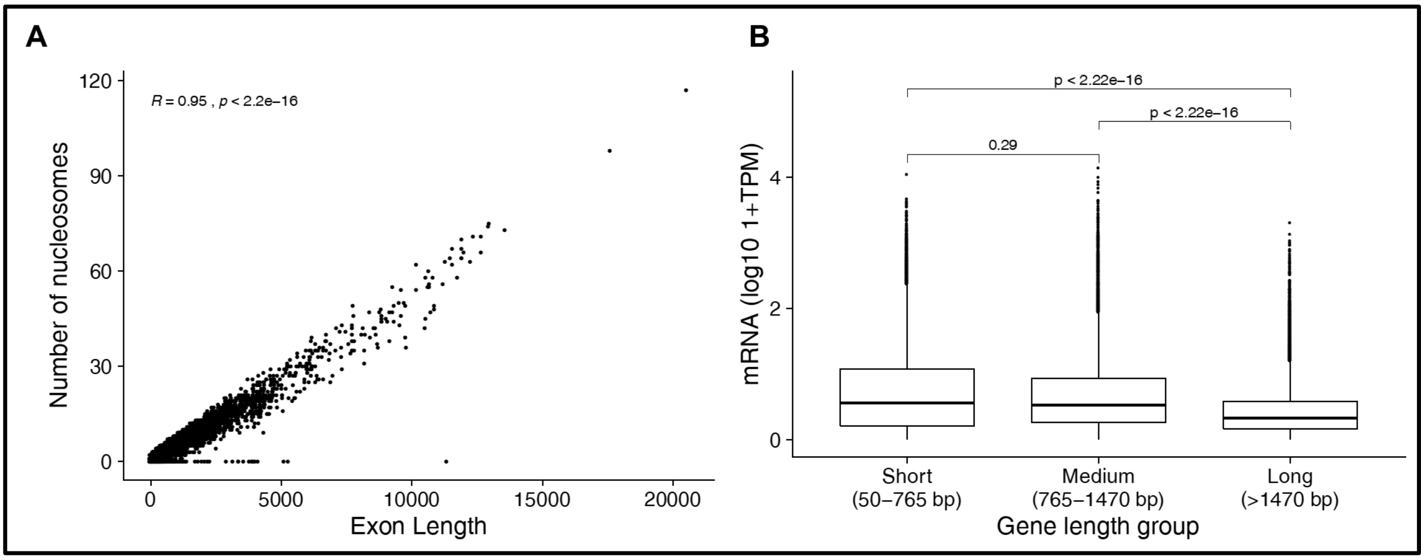


**Suppl. Fig. 10** A) Scatter plot shows the increasing number of nucleosomes (y-axis) proportional to the increasing exon length (x-axis). Pearson correlation value, and its statistical significance is shown embedded in the plot. B) Gene expression stratified according to different gene length groups.

### Suppl. Fig. 11

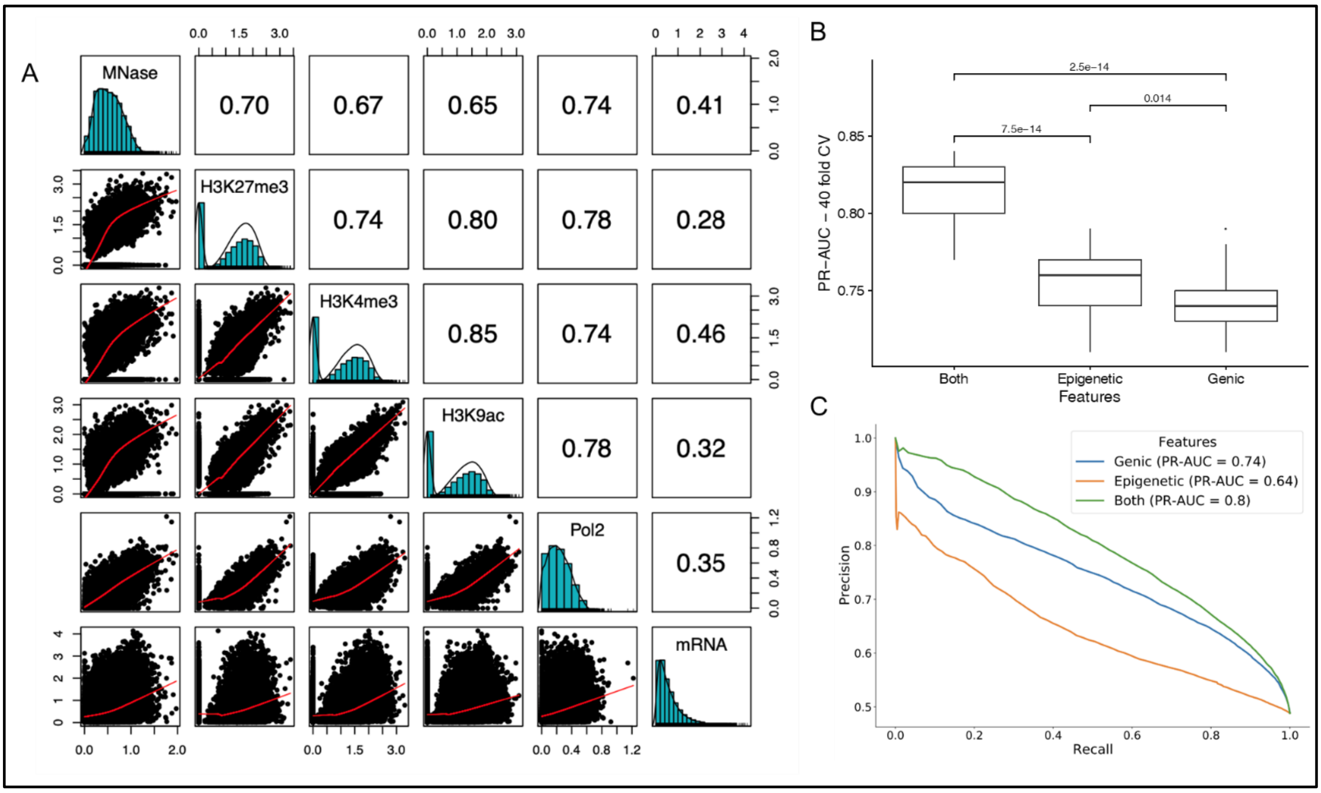


**Suppl. Fig. 11** A) The distribution plot of each epigenetic mark and mRNA are shown along the diagonal. The Pearson correlation coefficients (above the diagonal) are shown for the respective variables mentioned along the x- and y-axis of each box. The y-axis of scatter plots (below the diagonal) belongs to the variable mentioned along the horizontal line of that plot. mRNA is measured in TPM units. All values shown are log10 transformed with a pseudocount of 1. B) A box plot showing the distribution of PR-AUC values of all 40-fold CV (y-axis) for random forests model using different feature subsets. C) A precision recall curve showing the average precision and recall values from a 40-fold cross validated (CV) random forests algorithm employing different feature sets (colors) to classify gene expression as high or low using the length normalized epigenetic signals from TSS+300 bp window, as opposed to whole gene body signals in the main Figure 7.

### Suppl. Fig. 12

A

| Plastic gene group | Low | High |
|---|---|---|
| G1 | 2433 | 7041 |
| G2 | 5104 | 5391 |
| G3 | 5645 | 4153 |
| G4 | 6737 | 2490 |

B

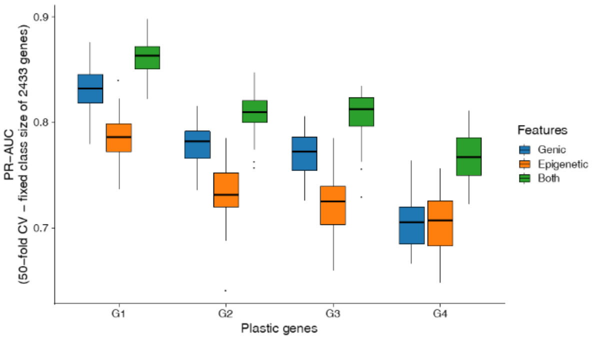


**Suppl. Fig. 12** A) The number of high and low expressed genes in each plastic gene group is shown. B) Box plot showing the distribution of a 50-fold cross-validation based PR-AUC of random forests models with different feature subsets. The number of genes in each plastic gene group were randomly subsampled to have 2433 genes in high and low expressed category. We chose 2433, as it is the lowest number of genes among the high/low expressed plastic genes.

### Suppl. Fig. 13

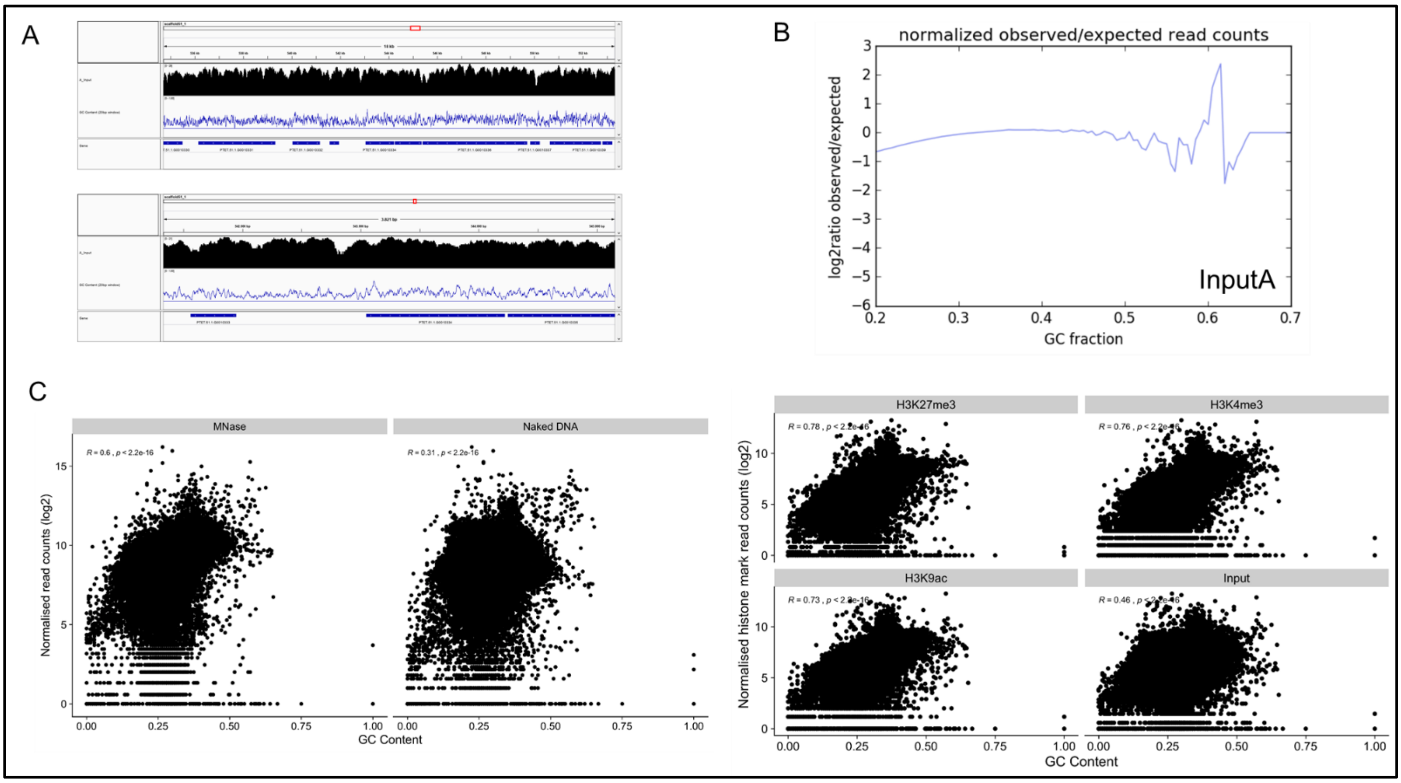


**Suppl. Fig. 13** A) Snapshots from Genome browser showing coverage track of ChIP input along scaffold51_1(black) and GC content of the underlying DNA sequence in 20 bp windows (blue line). Genes are shown in dark blue in the third row. The top panel shows the view in a 18kb window while the lower panel shows the zoomed view in a 3 kb window including an intergenic region. B) Correlation of reads from ChIP experiments to GC content. After binning the genome in 147 bp bins, the reads in each bin were counted, normalized and correlated against the GC content. B) The expected GC profile by counting the number of DNA fragments of a fixed size (by default is 300bp) per GC fraction is compared to the observed profile which is generated by counting the number of reads per GC fraction (the lower panel). C) Scatter plots of the GC content (x-axis) vs bin-length normalized read counts (y-axis; log2) of raw MNase, naked DNA (left), H3K27me3, H3K4me3, H3K9ac, and Input (right) measured in 147 bp bins of the genome. The Pearson correlation values are mentioned on the top left corner of the plots.
